# Chromatin arranges in chains of mesoscale domains with nanoscale functional topography independent of cohesin

**DOI:** 10.1101/566638

**Authors:** Ezequiel Miron, Roel Oldenkamp, Jill M. Brown, David M. S. Pinto, C. Shan Xu, Ana R. Faria, Haitham A. Shaban, James D. P. Rhodes, Cassandravictoria Innocent, Sara de Ornellas, Harald F. Hess, Veronica Buckle, Lothar Schermelleh

## Abstract

Three-dimensional (3D) chromatin organization plays a key role in regulating mammalian genome function, however many of its physical features at the single-cell level remain underexplored. Here we use 3D super-resolution and scanning electron microscopy to analyze structural and functional nuclear organization in somatic cells. We identify linked chromatin domains (CDs) composed of irregular ∼200-300-nm-wide aggregates of nucleosomes that can overlap with individual topologically associating domains and are distinct from a surrounding RNA-populated interchromatin region. High-content mapping uncovers confinement of cohesin and active histone modifications to surfaces and enrichment of repressive modifications towards the core of CDs in both hetero- and euchromatic regions. This nanoscale functional topography is temporarily relaxed in post-replicative chromatin, but remarkably persists after ablation of cohesin. Our findings establish CDs as physical and functional modules of mesoscale genome organization.

## INTRODUCTION

The genome in mammalian cell nuclei is hierarchically organized at various scales correlating with diverse genomic functions (*1*, *2*). At the base pair to kilobase pair level, DNA is wrapped around core histones to create nucleosomes (*3*). At the 100 Mb scale, entire chromosomes harbor distinct territories within the nucleus with transcriptionally active euchromatic and inactive heterochromatic segments tending to segregate into specific nuclear sub-regions (*4*). Chromatin organization at the mesoscale ranging from several kb to 100 Mb remains poorly understood (*5*). Advances in next-generation sequencing-based chromosome conformation capturing methods (3C/Hi-C) have revealed partitioning into several hundred kb to a few Mb-sized topologically associating domains (TADs) (*6*, *7*). TADs are genomic segments with higher intra-TAD contacts compared to inter-TAD contacts. At higher levels of organization, TADs group into ∼10-20 Mb genomic A- and B-compartments. A-compartments denote DNase I sensitive, transcriptionally active ‘open’ chromatin and B-compartments denote transcriptionally repressed, ‘closed’ chromatin (*6*). Compartments are generally correlated with euchromatin or heterochromatin, nuclear interior or lamina/nucleolar contacts for A- and B-compartments, respectively (*8*).

A key regulator in TAD organization is CCCTC-binding factor (CTCF), which binds to convergently oriented recognition sequences that flank TADs, and which define TAD boundaries at the linear genomic (1D) scale (*9*). Deletion of genomic TAD boundaries leads to aberrant transcriptional output (*10*). Co-occurrence of the ring-shaped cohesin complex at CTCF binding sites suggest a regulatory role for cohesin in shaping TAD structures, possibly through a loop extrusion mechanism (*11*–*16*). Accordingly, loss of cohesin function leads to an erasure of TAD signatures on Hi-C interaction maps (*17*).

Whereas the linear genomic size of TADs is defined, their spatiotemporal organization is underexplored. Single-cell Hi-C and more recent microscopy-based approaches using fluorescence *in situ* hybridization (FISH) of TAD-based regions have observed a high degree of stochasticity and heterogeneity in chromatin conformations in fixed cells (*18*–*24*). Despite such advances in imaging single TADs, it is still unclear whether TADs always form a single physical entity, how they relate to adjacent nuclear compartments and how active and silenced chromatin is distributed at TAD scales.

*In silico* simulations of chromatin as a melted polymer suggest that the charged properties of modified histone tails alter the compaction status of such chromatin domains (*25*). This model explains the increase in the physical size of epigenetically active domains compared to epigenetically inactive domains (*26*). Decreased protein accessibility with increasing chromatin density may allow size differences between small transcription factors (TFs) and large nuclear macro-complexes, such as the transcription machinery, to be exploited as a mechanism of transcriptional control (*27*).

Here we use quantitative 3D super-resolution imaging of both fixed and live samples, complemented by 3D electron microscopy of cryo-preserved samples, to establish chromatin domains (CDs) as TAD-sized irregularly-shaped dense nucleosomal aggregates in the size range of ∼200-300 nm. CDs are linked sequentially in a 3D curvilinear arrangement to form a distinct reticular network that is separated by a relatively wide RNA-filled interchromatin space. We apply a novel high-throughput image analysis pipeline for 3D-SIM data to spatially map functional markers, such as structural proteins, histone modifications and RNA polymerase II (RNAPII). We describe for the first time at physically relevant nanometer scale distinct volumes or zones of genome function in the context of CDs. This organization is independent of cohesin, establishing CDs as functional and physical modules of mesoscale genome organization in mouse and human somatic cells.

## RESULTS

### Chromatin arranges in coherently moving sub-micrometer domains, distinct from an RNA-occupied interchromatin compartment

The study of <1 Mb chromatin domain topologies in single cells *in situ* and *in vivo* has long been hindered, because they fall beyond the diffraction limit of light microscopy. Advances in super-resolution microscopy now put such structures within visual reach (*20*, *24*). 3D structured illumination microscopy (3D-SIM) enables fast multicolor 3D acquisition of whole cells with 8-fold higher volumetric resolution and strongly enhanced contrast compared to conventional fluorescence microscopy (*28*). Previous 3D-SIM studies resolved 4’,6-diamidino-2-phenylindole (DAPI)-stained chromatin in mammalian somatic cells as an intricate sponge-like structure, juxtaposed to an interchromatin compartment (IC) of no detectable chromatin that ends in channels leading to nuclear pores (Fig. 1A) (*29*–*31*). Using rigorously quality-controlled 3D-SIM (Methods, Fig. S1A) in mouse (C127) and human (HeLa) somatic cells, we observe chromatin as a chain-like reticular structure of distinct nodes or chromatin domains (CDs) of variable diameters in the range of a few 100 nm, hereafter referred to as CD chains. DNA staining with either DAPI or SYTOX Green both show the same sponge-like pattern (unlike DAPI, SYTOX Green exhibits no AT-base preference) (Figs. 1A and S1A). Control experiments in HeLa cells stably expressing histone H2B-GFP showed a near complete co-localization of H2B-GFP and DNA staining (Fig. S1B). Live cell 3D-SIM imaging (Methods) confirms the presence of a CD chain – IC arrangement *in vivo* (Fig. S1C) and unveiled a highly dynamic behavior of CD features over different observation periods and time intervals (Fig. S2, Movies S1-S3). Furthermore, quantitative flow field analysis (*32*) (Methods, Fig. S2C, Movie S4) reveals localized spatially coherent chromatin motion, i.e. fields with same motion direction. These properties are consistent with mechanically-coupled viscoelastic droplet-like domains (*32*, *33*), or ‘blobs’.

**Fig 1.**
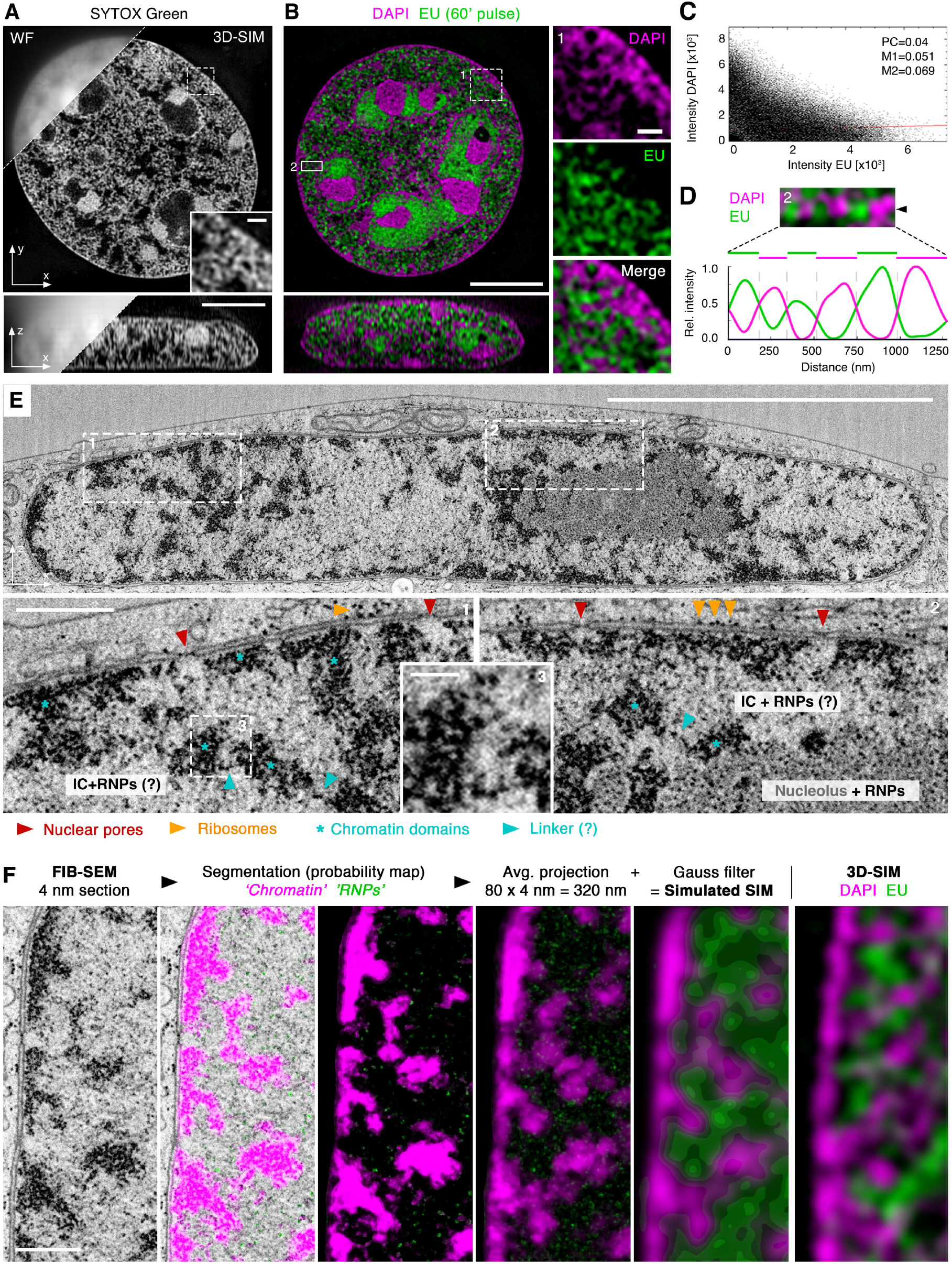
Chromatin folds into 3D nanodomains of nucleosomal aggregates distinct from an RNA-occupied interchromatin network. (**A**) In contrast to conventional widefield (WF, top left), super-resolution 3D-SIM of fixed mouse C127 mammary epithelial cells resolves SYTOX Green stained chromatin as a sponge-like reticular structure throughout the nuclear volume. Chromatin is separated by an unstained interchromatin (IC) space. Top panel shows a lateral, bottom panel an orthogonal cross section through the 3D image stack. Scale bars: 5 *μ*m and 0.5 *μ*m (inset). See also Fig. S1. (**B**) IC juxtaposed to DAPI-stained chromatin is largely occupied by bulk RNAs stained with EU (60 min pulse). Right, inset magnifications of region 1, and below, orthogonal cross section. Scale bars: 5 *μ*m and 0.5 *μ*m (inset). (**C**) Pixel-based colocalization analysis scatter plot confirms anti-correlation of DAPI and EU signals. PC: Pearson correlation coefficient; M1/M2: Manders’ correlation coefficients DAPI vs. EU and EU vs. DAPI, respectively. (**D**) Line profile of signal intensity for both DAPI and EU from a representative euchromatic region 2 in panel (B), highlighting the mutually exclusive volumes occupied by DNA and RNA that are in the size range of 200-300 nm. (**E**) Single FIB-SEM section of a cryo-fixed HeLa cell, stained with osmium and uranyl acetate under cryo-conditions before resin embedding and imaging. Consecutive 4 nm section were cut by a focused ion beam (FIB) in orthogonal direction to growth surface and scanning electron microscopy (SEM) was recorded with 4 nm pixel size. The nuclear volume indicates G1 (or early S phase) stage. Scale bar: 5 *μ*m. Below: inset magnifications of two indicated boxes with postulated features annotated. Red arrowheads indicate sites of nuclear pore complexes (NPCs) within the double membrane of the nuclear envelope. Blue asterisks indicate 200-300 nm wide aggregates of 10-20 nm diameter sized dark dots, likely nucleosomes. Orange arrowheads indicate ribosomes at the outer nuclear membrane. Blue arrowheads indicate putative linker segments of single nucleosomes. Scale bars: 200 nm inset 1 and 2 and 100 nm inset 3. (**F**) Columns 1-3: lateral cross section of the dataset shown in panel E followed by machine learning assisted segmentation of chromatin and putative RNPs. Columns 4-6: Average intensity projection of 80 consecutive segmented FIB-SEM sections covering a 320 nm z-range followed by 3D-Gauss filtering to match 3D-SIM’s spatial resolution, closely resembles 3D-SIM micrograph at the same magnification. Scale bar: 0.5 *μ*m. See also Fig. S2.

We next sought to better understand the content of the relatively wide IC space separating these domains. Considering RNA transcripts as a prime candidate, we performed an extended 5-ethenyluridine (EU) pulse labelling to detect all transcribed RNAs (*34*). We observed a striking enrichment of bulk RNA in the IC space including nucleoli and nuclear pore channels, and almost mutual exclusion with DAPI-stained chromatin (Fig. 1B, C). Line profiles through a representative euchromatic region of the nuclear interior highlight DNA domains in the size range of 200-300 nm alternating with the RNA-containing IC space of similar dimension (Fig. 1D).

The resolution limit of our 3D-SIM imaging approach is ∼115 nm at 488 nm excitation (Fig. S1A). To test if the observed nodal pattern of chromatin derives from structures at the molecular scale, we made use of high-resolution focused ion beam scanning electron microscopy (FIB-SEM). This approach allows imaging of entire nuclear volumes at 4 nm isotropic resolution (*35*). HeLa cells were fixed and stained under cryo-conditions, before resin-embedding, enhancing the contrast of nucleoproteins and ensuring the best possible structural preservation (*36*). Assessment of FIB-SEM of HeLa (G1 and G2) and U2-OS cell nuclei (Fig. 1E and S3A, B) allows identification of distinct CDs as densely packed aggregates of hundreds of highly contrasted ∼10 nm sized dots (likely nucleosomes) typically connected by linker filaments (Fig. 1E, Movie S5). Interestingly, chromatin at the nuclear periphery is seen as an inseparable melt, creating a homogeneous, almost continuous chromatin layer, except for NPC holes (Fig. S3C). Using a machine learning approach (*37*), we segmented CDs of aggregated nucleosomes distinct from nucleolar and IC regions (Movie S6, Figs. 1F, columns 1-3 and S3D-F). The remaining sites marked solitary 10-20 nm sized nuclear particles (likely RNA-protein complexes such as RNPs or spliceosomes) within the IC. Projection and convolution with a Gaussian function of chromatin and RNPs segmentation probability maps enabled us to generated a close mimic of 3D-SIM micrographs of DNA/RNA stained cells (Fig. 1F, columns 4-6). Quantitative measurements of CD widths along the nuclear envelop yielded an average extend of ∼200 ±60 nm. Throughout the nuclear volume, the bulk of CDs have a dimension of between 200-300 nm (Fig. S3D). Smaller CDs (down to 100 nm) and larger CDs (up to > 500 nm) are also apparent, but less frequent (Movie S5).

We conclude the bulk of nuclear RNA and DNA occupy mutually exclusive volumes. Both, 3D-SIM and FIB-SEM independently identify chromatin as a convoluted, reticular structure composed of chains of sub-micrometer CDs.

### CDs colocalize with TAD sequences and follow a curvilinear arrangement

Having established the arrangement of chromatin into spatially confined CDs, we used DNA FISH detection to place the observed 3D chromatin landscape in the context of physiological TADs as defined by Hi-C experiments. To avoid harsh denaturation and disruption of chromatin structures below ∼1-Mb-size levels (*38*), we have implemented a non-denaturing FISH method, termed resolution after single-stranded exonuclease resection (RASER)-FISH (*39*, *40*). We applied RASER-FISH to study the topology of a previously characterized 0.7 Mb TAD ‘H’ located on the mouse X chromosome (*22*) (Fig. 2A and S4A). Quantitative analysis of 21 TAD H RASER-FISH signals revealed an average core diameter of ∼330 nm (Fig. S4B). Although TADs in single cells are mostly globular they are not uniform and some extend in at least one dimension to a maximum of ∼500 nm (Fig. S4B). Importantly, the spatial boundary, or rim of TAD FISH signals coincides with the rim of underlying CDs (Fig. 2A, B). Furthermore, multi-color RASER-FISH of neighboring genomic domains spanning the *α-globin* locus revealed CDs arranged in a curvilinear path (Fig. 2C). Each genomic segment forms a discrete 3D nanodomain along a convoluted chain irrespective of their relative transcriptional activity (Fig. S4C).

**Fig 2.**
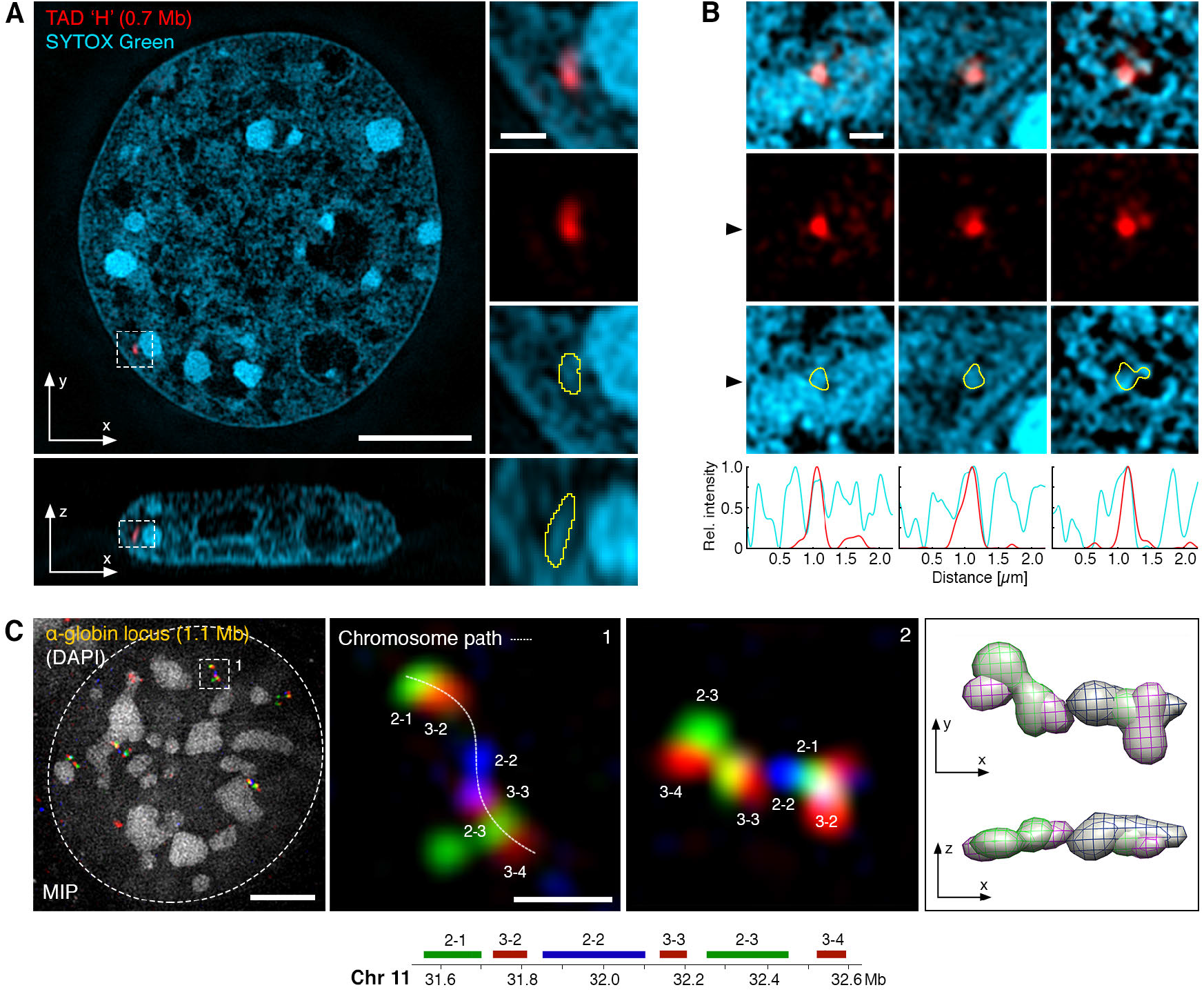
Spatial relationship between CD chain and TADs. (**A**) RASER-FISH detection of a 0.7 Mb X chromosomal TAD ‘H’ (red; Hi-C map shown in Fig. S4A) and SYTOX Green whole chromatin labelling (cyan) in a mouse C127 cell imaged by 3D-SIM. Right, inset magnifications and below, orthogonal cross section. Yellow outlines of the FISH signal highlight ta close overlap with underlying CD chain nodes. (**B**) Further representative examples of TAD ‘H’ FISH signals from separate cells outlined against chromatin. Below, line profile of signal intensity for both TAD H (red) and SYTOX Green (cyan) calculated from a horizontal cross section at the black arrowhead. The spatial boundary of TAD H correlates strongly with CD boundaries. See also Fig. S4A, B. (**C**) Multicolor RASER-FISH labelling of sequential TADs across the α-globin gene locus detected in mouse C127 cells. Maximum intensity projections (MIP) of 3D-SIM micrographs show the genomic order of the probes is preserved as a curvilinear chain of discrete CDs in space. Note that DAPI has only low affinity for single-stranded DNA and is only used here to indicate the nuclear outline. A second example region from a different cell is shown on the right as MIP and 3D rendering (with views from different angles). Scale bars: 5 *μ*m and 0.5 *μ*m (insets). See also Fig. S4C.

We conclude that chromatin in somatic interphase cells arranges as a convoluted curvilinear chain partitioned into discrete non-rigid shaped (globular or extended) CDs. These can harbor TADs and are separated from a DNA-depleted, RNA-occupied interchromatin compartment.

### Functional marker distribution reveals 3D zonation at CD scales

The functional output of DNA is influenced by a broad repertoire of protein markers such as histone modifications, architectural proteins, transcription, replication, and chromatin remodeling enzymes. In order to understand how key functional markers mapped against the nuclear landscape at the nanoscale we have devised a pipeline for automated high-content image analysis for the mining of spatial information from thousands of multicolor-labelled 3D-SIM nuclear volume datasets (for details see Fig. S5 and Methods). It first divides the chromatin signal into seven classes based on voxel intensity (*31*), which serve as a proxy for positioning other stainings relative to CD features (Figs. 3A and S5E). The lowest intensity region, class 1 denotes the DNA depleted IC, classes 2 and 3 denote the outer rim or surfaces of CDs, previously termed perichromatin (PC; Fakan and van Driel, 2007), and classes 4-7 constitute the more interior core region of the CDs, with classes 6 and 7 denoting primarily constitutive heterochromatin regions. We characterized the 3D epigenome topography using a wide range of markers (16 in total) associated with specific genome functions (nascent RNA, RNA-associated proteins, chromatin-associated proteins and histone post-translational modifications (PTMs; Figs. 3B and S6A) in over 400 mouse C127 cells in cell cycle stage G1 (Table S1).

**Fig 3.**
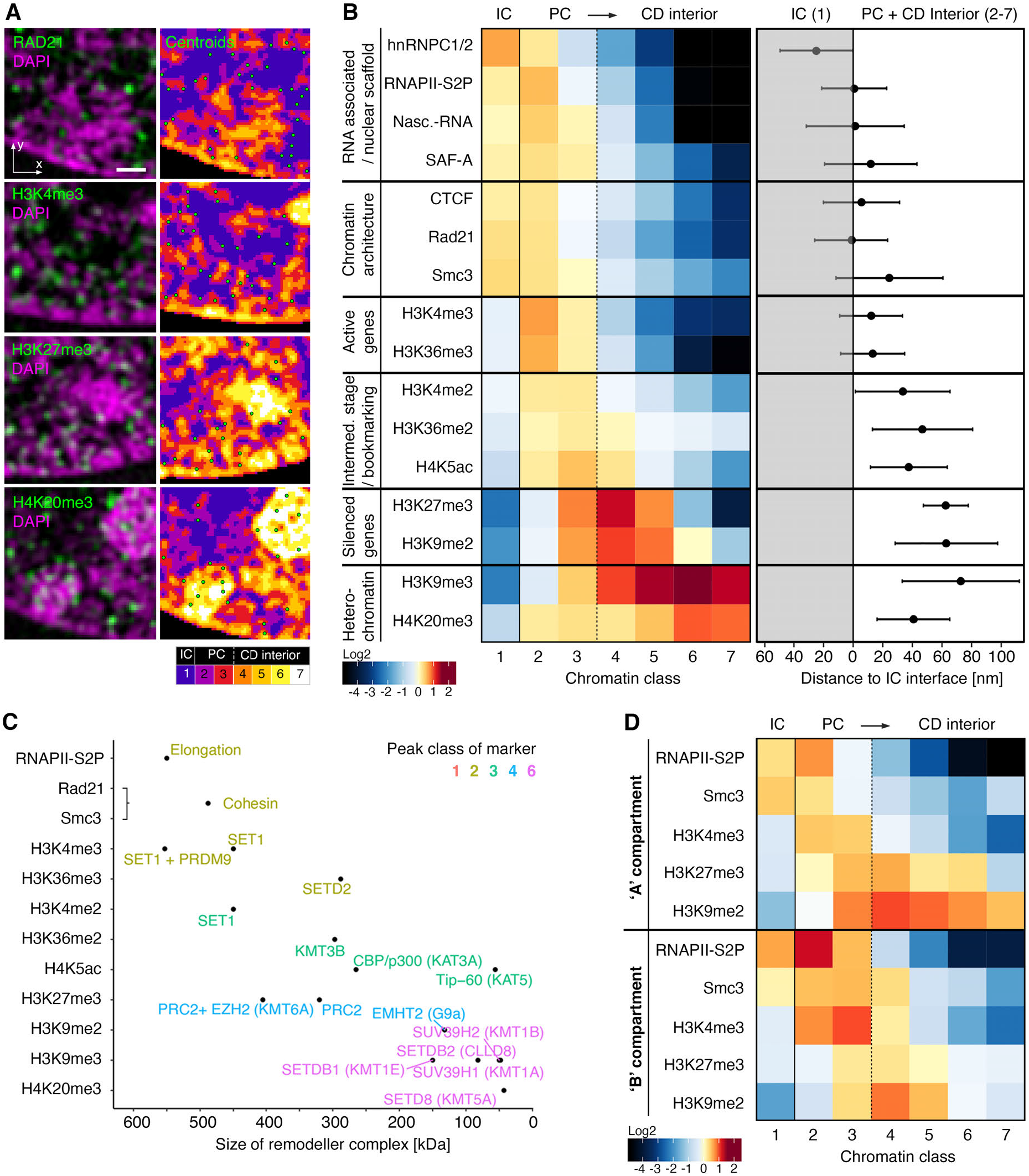
Functional marker distribution in somatic cells reveals nanoscale 3D zonation correlated with protein complex size. (**A**) Left, representative sub-regions of single z-planes of 3D-SIM image stacks of mouse C127 cells in G1comparing the spatial localization of different markers with respect to chromatin. Right, marker centroid coordinates are depicted as green dots with a black outline. Note that centroids are determined with subvoxel accuracy and may be closer to adjacent z-sections, and thus not be displayed in the chosen plane. G1 cell cycle stage was confirmed by the absence of EdU incorporation and relatively small nuclear volume. Scale bar: 0.5 *μ*m. (**B**) Left: Heatmap of enrichment or depletion of IF signals relative to a random distribution (plotted in log2-fold change) for each chromatin class for each marker. Right: Distance to the IC interface for all markers. The mean distance and 95% confidence interval of the population are shown. At least 20 cells in 2 replicates were recorded for each marker, in total *N* = 433 cells. (C) Molecular weights of chromatin complexes corresponding to each marker. Scatterplot highlights an inverted correlation of size and preferential marker distribution (peak chromatin class color-coded). MW for cohesin was calculated from Smc1, Smc3, Rad21 and SA1/2. (D) Heatmap of relative IF signal distributions of representative markers in nuclear sub-volumes harboring mainly heterochromatin characteristic of B compartments (the segmented perinuclear rim and chromocenter regions), or euchromatin characteristic of A compartments (the remaining volume). Both compartments show a relatively conserved zonation pattern despite the different composition of chromatin in either region (Fig. S5D). Total number of cells *N* = 170.

The RNA processing/interacting factors hnRNPC1/2, SAF-A (hnRNPU), RNAP-S2P and nascent RNA transcripts are noted as enriched in class 1 (the IC) and the perichromatin classes 2 and 3, depleted in classes 4 and 5, and are virtually absent in the highest classes 6 and 7 (Fig. 3B, top left). A recent study has shown the same pattern to be true for SC35, a protein involved in transcriptional splicing speckles (*42*). Interestingly, those factors known to interact with chromatin, RNAPII-S2P, nascent RNA and SAF-A, show a peak of enrichment in class 2, while the sole factor that only binds RNA, hnRNPC1/2, shows the strongest enrichment in class 1. Accordingly, average nearest-distance measurements to segmented chromatin surface result in a mean distance location of hnRNPC1/2 outside, while the other three markers center right at the border-line (Fig. 3B, top right). These results are in accordance with early immuno-EM observations that active transcription is confined to the PC (*41*). They are also in line with the reported interaction of SAF-A with chromatin-associated RNAs (*43*) supporting the concept of a physical 3D interface between an RNA environment and de-condensed outer CD fringes (Fig. 1B).

Unlike RNA-interacting proteins, the spatial distribution of segmented histone PTMs, which are inherently chromatin-associated, show a marked depletion in the lowest intensity class 1, further validating such annotated voxels as a true interchromatin compartment (IC). Importantly, we observe a biased enrichment for both H3K4me3, typically located at promoters of active genes, and H3K36me3, typically enriched along the body of transcribed genes (Fig. 3B, center left), in the lowest chromatin classes 2 and 3 marking the PC, with peak enrichment in class 2 similar to RNAPII-S2P and nascent transcripts. These markers are rarely found in the higher intensity classes 4 and 5, typically associated with the core of CDs, and are almost absent in the classes 6 and 7 (Fig. 3B, center left). Their corresponding di-methylated forms (H3K4me2 and H3K36me2), as well as acetylated H4K5, which has been implicated in epigenetic bookmarking (*44*) also show a notable albeit less distinct enrichment in the lower class range (peak in class 3) and depletion in higher classes.

In contrast, repressive histone PTMs generally show a broader distribution that is shifted towards the higher intensity classes (Fig. 3B, bottom). The H3K27me3 mark, which is typically deposited along Polycomb-mediated silenced genes (*45*) shows an enrichment in the interior classes (peaking at class 4). H3K9me3, a marker for transcriptionally inactive facultative and constitutive heterochromatin (*45*), is found enriched towards the CD interior and in heterochromatin classes (3-7), but is most abundant in the higher classes with a peak at class 6. H4K20me3, which has been implicated in silencing repetitive DNA (e.g. in chromocenters of mouse cells) and transposons (*46*), is most strongly enriched in classes 6 and 7, confirming these classes as constitutive heterochromatin. H3K9me2 shows a notably less shifted distribution than H3K9me3 peaking at the interior classes class 4 similarly to H3K27me3.

The observed differential enrichment of functional markers can also be considered in light of the specific enzyme complexes responsible for their deposition. Indeed, there is a remarkable inverse correlation, with smaller complex sizes correlating to peak enrichment in higher chromatin classes of the corresponding marker (Fig. 3C).This supports the hypothesis that chromatin density acts as a higher-level regulator of genome function by hindering physical accessibility of larger complexes to substrate chromatin (*27*). Analyzing a subset of markers in human colon (HCT116) and cervical (HeLa) cancer cell lines in G1 stage, we found highly similar differential spatial distributions, confirming the universal nature of the observed zonation (Fig. S6B, C).

It has been postulated that TADs may be wholly accessible or inaccessible depending on whether they form part of the A or B compartment respectively, as denoted by Hi-C experiments. To test this, we sub-divided the nuclear volume into lamina-associated chromatin plus chromocenter regions, or the residual nuclear volume, as proxies for general B and A compartmentalization, respectively. We find the nanoscale zonation true for both euchromatic (‘A’) regions as well as heterochromatic (‘B’) regions (Figs. 3D and S6D).Thus, both large-scale compartments harbor transcriptionally active and inactive sequences with nanoscale distributions, albeit at different ratios (*47*) (Fig. S6E).

In conclusion, our high-content 3D super-resolution mapping approach highlights an ordered zonation of epigenetic markers, forming a nanoscale functional chromatin topography. The propensity of different functions (e.g., transcription silencing) to occupy different chromatin classes is independent of whether that chromatin is found in euchromatic or heterochromatic regions. Markers of active transcription and architectural proteins are confined to the PC (and CD linker regions), while silencing modifications are enriched towards the CD interior. The IC harbors most RNAs and RNA interacting proteins. Finally, we suggest the packing of nucleosomes into mesoscale CDs may lead to the physical exclusion (or hindrance) of chromatin-associated complexes according to their size, thus providing a possible explanation for the ordered functional zonation as a consequence of physical accessibility (*27*).

### The local CD topography is temporarily relaxed during DNA replication

DNA polymerase is a large complex and is known to begin replication in a characteristic nuclear pattern of replication foci (*48*). Such observations are in agreement with our finding of larger complexes being enriched at lower chromatin classes. However, DNA replication must eventually process the entire genome, irrespective of the activity state at any given locus. We hypothesize that every chromatin locus should, over time, be locally decompacted in the process of replication. To test this at the nanometer scale, we identified S-phase cells by short EdU pulse labelling and grouped them into early, mid and late S-phase stages. Sub-dividing the nucleus in this manner reveals a strong local de-compaction at sites of ongoing replication as revealed by chromatin classification in the local vicinity of EdU foci (Fig. 4A). The disruption is confined to the local activity of the replication machinery as no significant changes could be seen when analyzing the entire nuclear volume (Fig. S7A). Local de-compaction is dramatically highlighted by tunnel-like voids becoming apparent in otherwise dense chromocenters in late S-phase (Fig. 4A, right). This is in agreement with a recent study of DNA synthesis at constitutive heterochromatin (*49*). Accordingly, EdU labelled replication foci show an enrichment in the lower chromatin classes, irrespective of S-phase stage (Fig. 4B).

**Fig 4.**
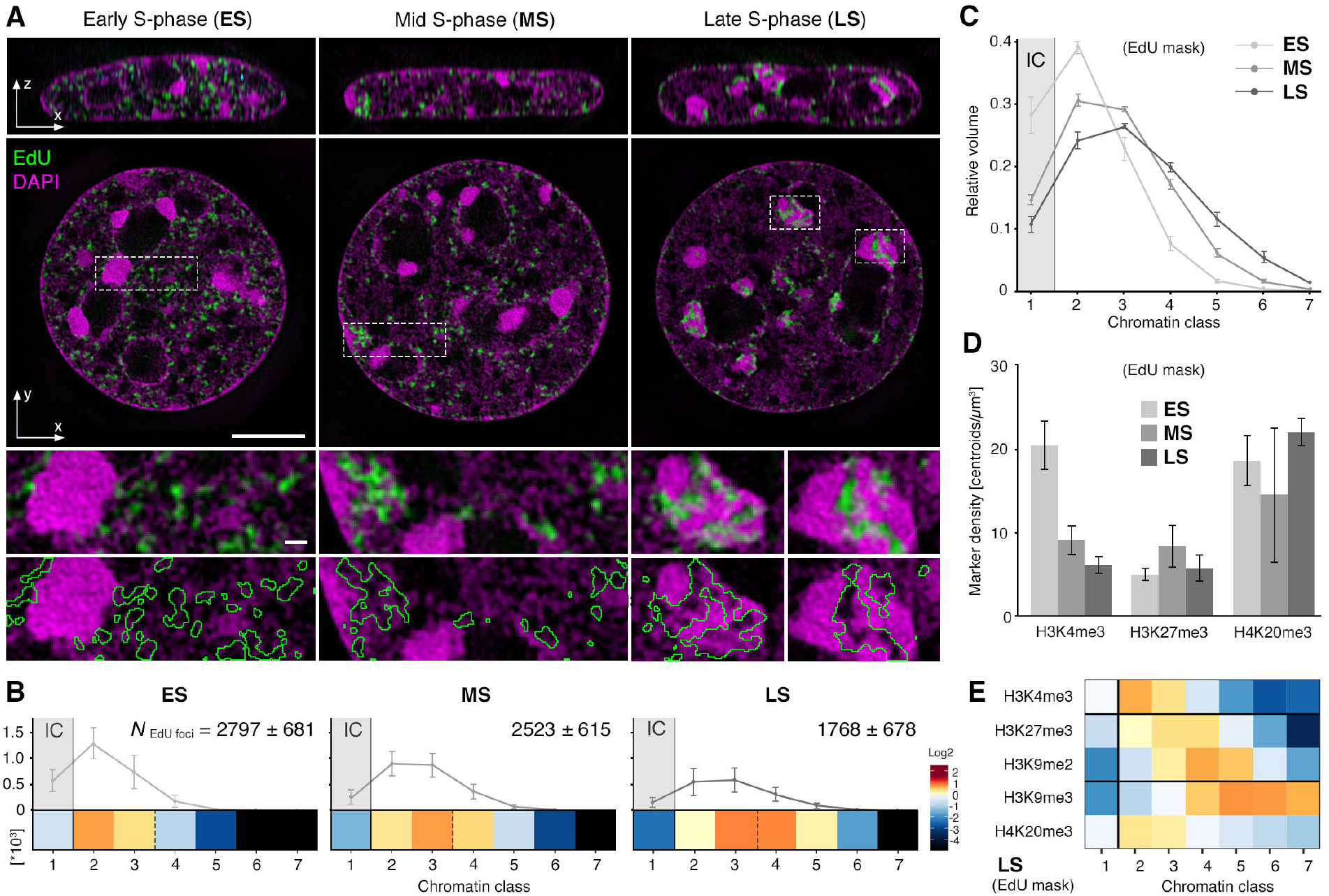
Effect of active replication on chromatin composition.

(**A**) Orthogonal and lateral cross sections of representative 3D-SIM datasets showing the chromatin landscape with the different patterns of replication foci (green) during S-phase progression from early, mid and late S-phase stages (left to right). Chromatin is stained with DAPI and replication is labelled using 15 min of EdU incorporation. Local replication regions were masked based on the EdU signal and outlined in green. Scale bar: 5 *μ*m and 0.5 *μ*m (insets). (**B**) Quantification of the absolute number of detected EdU foci in each of the different chromatin classes. The total number of EdU foci per cell is indicated as mean ± SD. Per S-phase stage, a heatmap of enrichment or depletion of EdU signals relative to a random distribution (plotted in log2-fold change) is depicted. (**C**) Relative chromatin class distribution within EdU submask volumes in different S-phase stage. The mean ratios ± SD are depicted. (**D**) Foci number per *μ*m^3^ of EdU submask volume for the depicted histone modification in each condition, per S-phase stage. Mean ± SD are depicted. Number of cells: ES = 225, MS =227, LS =230. (**E**) Heatmap of statistical distribution of selected marker in EdU masked regions of late S phase cells highlights a relaxation of repressive marker confinement to CD chain interior compared to G1 shown in Fig. 3B. See also Fig. S7 for a complete list of marker distributions in all cell cycle stages.

When quantifying chromatin classes in replicating sub-regions, we observed, as expected, a shift towards denser chromatin classes when progressing from early, to mid, to late S-phase (Fig. 4C). Accordingly, the highest abundance of the active histone PTM H3K4me3 is in early S phase, as compared to the repressive PTM H3K27me3, which is most enriched in mid S phase (Figs. 4D and S7B). Constitutive heterochromatin marks, in particular H4K20me3 become less enriched in denser chromatin classes during late S-phase (Figs. 4 E and S7B) with the replication-driven opening of otherwise condensed chromatin.

Our data shows that active DNA replication can locally disrupt the physical organization of chromatin and consequently its functional zonation.

### Cohesin function is dispensable for 3D chromatin structure and functional zonation

While DNA polymerase is able to cause local rearrangements in S phase, other proteins are known to regulate chromatin structure globally throughout interphase. The cohesin complex is able to delineate the genomic boundaries of TADs (*50*) by a loop extrusion mechanism towards convergent CTCF binding sites (*16*). In our analysis, the distributions of CTCF and two sub-units of the cohesin complex, Smc3 and Rad21, were characterized by distinct enrichment in classes 1-2 and depletion in all higher classes (Fig. 3B left). They also showed a distance profile centering at the 3D-segmented CD chain-IC interface (Fig. 3B right). Cohesin degradation can erase TADs in Hi-C maps (*17*), but does not erase globular domains in FISH images (*20*). To address the role of cohesin on the global chromatin organization and the spatial zonation of the epigenome, we performed our analyses in human HCT116 cells with an auxin-inducible protease degradation system to ablate the RAD21 subunit of the cohesin complex (*51*) (Fig. 5A). Upon addition of both doxycycline and auxin, RAD21 can be depleted with a half-life of 17 min (*51*) and completely ablated in 2 h (Fig. S8A, B). We hypothesized that if the epigenomic zonation were governed by sequencespecific CTCF-guided cohesin activity, the 3D epigenome should be disrupted, while if this zonation were governed by non-sequence specific biophysical constraints it should not change after cohesin depletion. Strikingly we see no changes in the distribution of representative histone PTMs after 6 h of cohesin depletion (Figs. 5B and S8C).

**Fig 5.**
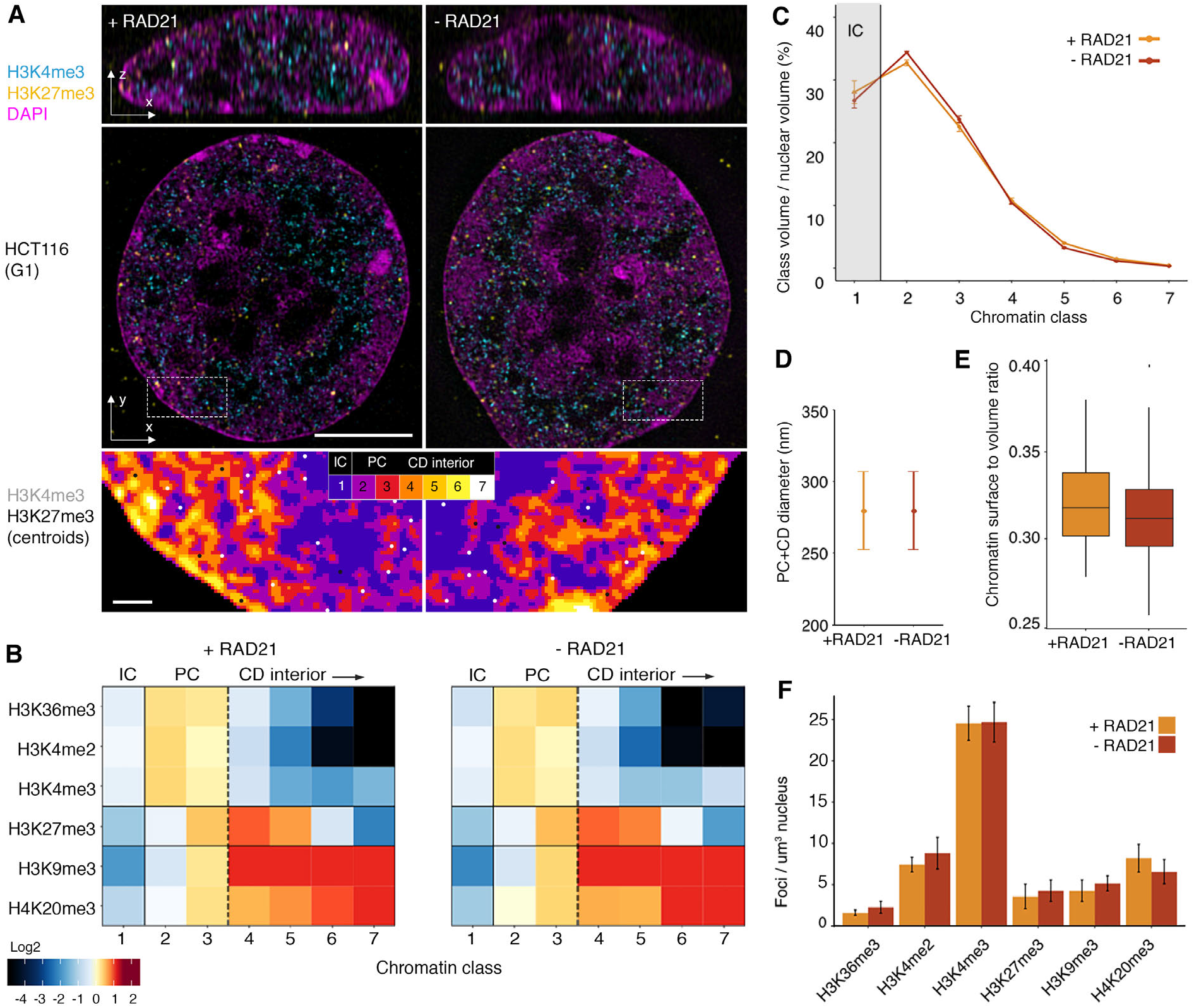
Epigenome zonation persists after cohesin ablation. (**A**) Representative single z-planes of 3D-SIM images comparing the chromatin landscape in human HCT116 in the presence of cohesin, and without cohesin. Control cells were only treated with auxin for 6 h (+RAD21, left) or RAD21 ablated cells after 16 h doxycycline and 6 h auxin treatment (-RAD21, right). Insets show the segmented topography with the spatial distribution of H3K4me3 and H3K27me3 represented as white and black dots respectively. Scale bars: 5 *μ*m and 0.5 *μ*m (insets). See also Fig. S8A, B. (**B**) Heatmap of enrichment or depletion of IF signals from a representative subset of histone modifications. Respective profiles for individual markers with error bars show no significant change for any distribution See also Fig. S8C. (**C**) Proportion of nuclear volume for each class, showing no significant change after ablation of cohesin function (mean ±95% CI). (**D-E**) Quantification of the PC+CD dimensions (D, ±95% CI, see Methods) and IC-PC surface area to PC+CD volume ratio (E, ±SD) shows no significant change between RAD21 positive or depleted conditions. (**F**) Quantification of the mean number of foci detected for each condition shows no significant change between RAD21 positive or depleted conditions (±SD). Total number of cells: + RAD21= 80; - RAD21= 104.

Furthermore, quantitative analysis of HCT116 RAD21-mAID cells 6 h after induction does not show any observable change in the CD chain structure (Fig. 5C-E), nor are there changes in the number of functional histone PTMs on this landscape (Fig. 5F). The CD chain diameter and the ratio of segmented chromatin surface area relative to its volume does not change in the absence of cohesin, suggesting that cohesin is dispensable for these structures. This phenomenon has recently been observed to persist over longer periods of depletion (*42*).

Our findings demonstrate that loss of cohesin, and thereby of TADs (as defined by Hi-C), has neither an observable effect on maintaining CD structures, nor on the epigenomic zonation relative to CDs in single somatic cells. We conclude that spatial functional zonation is regulated by the physics of the system independent of cohesin function, and therefore not defined by sequence-specific processes.

## DISCUSSION

By applying a rigorous quantitative high-content super-resolution 3D mapping approach of a wide range of functional markers, complemented by FIB-SEM, we establish that chromatin in somatic cells arranges in curvilinear chains of CDs of ∼200-300 nm diameter, forming physical modules that subdivide the mammalian genome into functional volumes (Fig. 6A). CDs can harbor individual TADs in single cells and are spatially juxtaposed to and mutually excluded from an RNA-rich interchromatin compartment (IC). The CD chain-IC organization features enrichment of diverse genome functions in different volumes, at the nanometer scale. Active transcription and architectural proteins are confined to the CD chain-IC interface and silencing marks are enriched towards the core of the nucleosomal aggregates (Fig. 6B).

**Fig 6.**
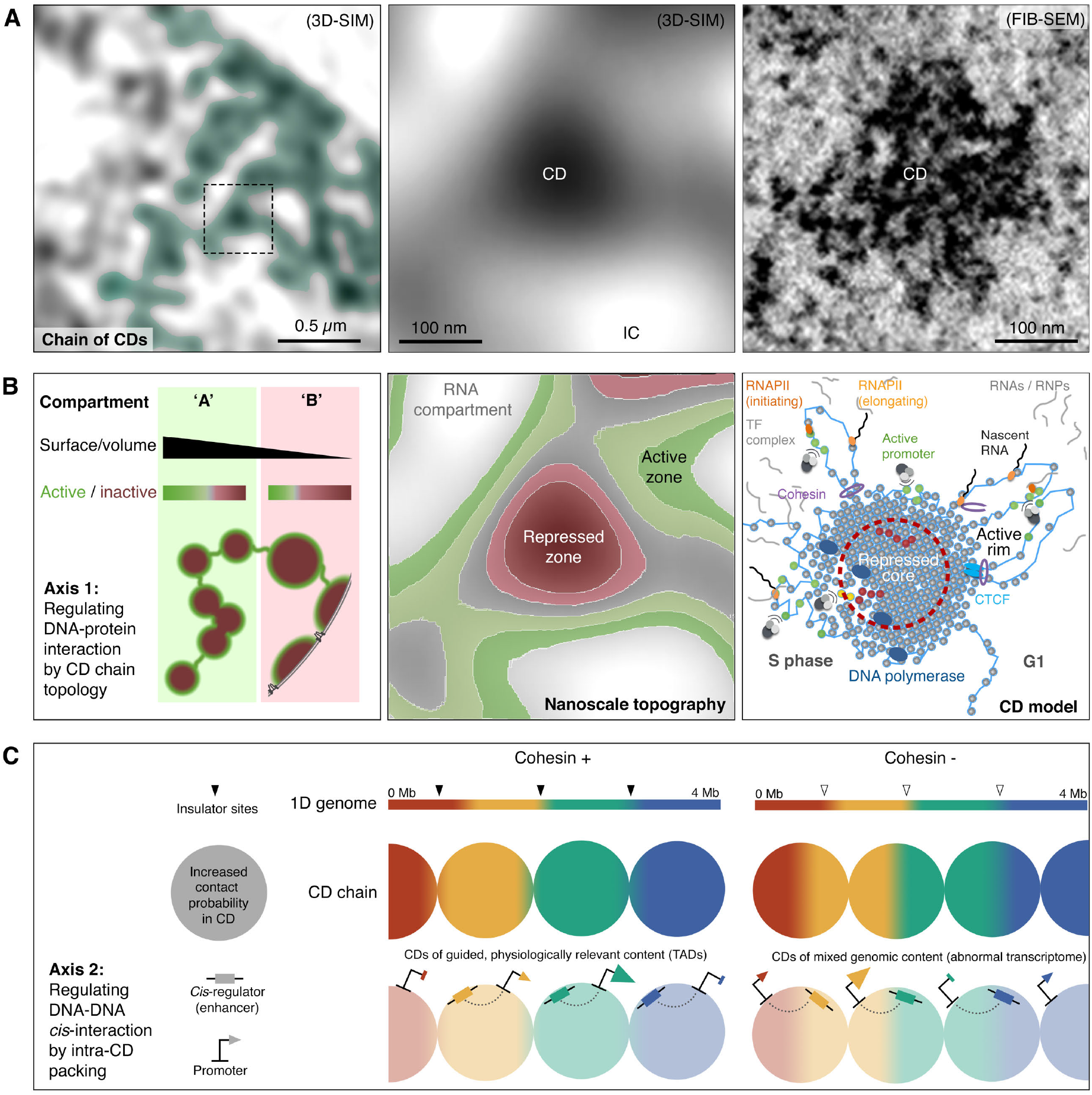
Model of the interphase nucleus as a chain of CDs from the mesoscale to the nanoscale. (**A**) Left: Super-resolved interphase chromatin stained with DAPI in a C127 cell, with an overlaid representation of a putative curvilinear chain of CDs. Center: Zoom-in to an individual CD, indicated by the black dashed box in the left panel, imaged by 3D-SIM. Right: an equivalently-sized individual CD imaged by FIB-SEM. (**B**) Model of functional zonation at each scale or resolution for images in panel A. Left: The ratio of active surface area to volume (SA/V) for DNA-protein interactions is the first regulatory axis of CDs over genome activity. Differing ratios of SA/V in either A or B compartments reflects changes in CD chains, and gives rise to generally active or inactive compartments. In this cartoon the ‘B’ compartment is adjacent to the lamina, represented by a black line interrupted by nuclear pores. Center: Preferential zonation of various functions occurs at the nm scale, relevant to single CDs. Right: The nanoscale zonation can be modelled by the radial position of histones with diverse PTMs and protein complexes. Almost no active markers are detected beyond ∼60 nm from the PC-IC interface towards the CD core, such that any silenced promoter (red spots) at this “depth” (within red dashed circle) may remain inactive by being inaccessible to the transcriptional machinery rather than through active deposition of repressive histone PTMs. Nevertheless, in S phase the local decompaction caused by DNA polymerase offers the chance for certain loci to escape inactivation, by being released from their inaccessible state (yellow spots). (**C**) Supporting appropriate DNA-DNA interactions is the second regulatory axis of CDs over genome function. The presence of cohesin links the 3D structure to the correct 1D genomic context, leading to physiologically relevant interactions observed as “TADs” in population Hi-C maps (left). Folding of the genome into biophysically defined CDs in the absence of cohesin leads to CD structures without physiological relevance (multiple colors mixed over genomic insulator boundaries, right).

### CDs provide an axis of control over DNA-protein interactions

We demonstrate for the first time that nanoscale zonation occurs in both euchromatic and heterochromatic compartments of the nucleus, deviating from previous models of wholly active ‘open’ vs inactive ‘closed’ domains. Based on our data, we postulate that genome segments rich in active sites display a topology of multiple smaller or extended volumes and a higher surface area. Segments depleted in active sites, in contrast, form fewer more globular CDs or melts along the nuclear/nucleolar periphery with reduced surface areas (Fig. 6B, left). The large fraction of non-coding DNA present in the genome despite the associated energetic costs in its accumulation and replication has always been puzzling. However, the potential of CDs to use excess DNA in nucleosome aggregates as regulation by means of accessibility may illuminate a *raison d’être* for much of the genome’s non-coding DNA.

CDs positioned at the nuclear envelope have less active interface due to the lamina blocking one flank. Observations from FIB-SEM reveal that CDs associated with the lamina, that are likely equivalent to lamina associated domains (LADs; Kind et al., 2013), also lack inter-CD separation by single nucleosome linkers (a feature of CDs in the nuclear interior). Instead, they are form a contiguous melt underneath the nuclear lamina. This conformation further reduces the available surface to just the nucleoplasmic flank (with the exception of channels reaching NPCs). Our model predicts this to favor silencing, which is true for LADs.

The described macro-scale organization can be spatially inverted while maintaining normal physiological function in the nuclei of nocturnal mammalian retina cells (*53*). A model where regulation occurs at the nanometer scale is independent of the nuclear position of the CD at scales of tens of microns, as long as the local CD surface to volume ratio can be controlled. This provides a mechanism for how tissue-specific transcriptional programs can be established despite large variation in the spatial arrangements of macro compartments between single cells (*54*), even in totally inverted architectures.

The ratio of active CD chain-IC interface area to CD chain volume should regulate genome activity by controlling the accessibility of nuclear machineries to their substrate, chromatin, although it is possible for some (e.g. DNA polymerase, Fig. 6B, right) to overcome these restrictions, reshaping the CD chain. We observe that zonation correlates with the size of chromatin-associated complexes, supporting *in silico* models. In previous modelling, accessibility of transcription factors (TF) is directly correlated to their molecular dimension, suggesting that binding of large TF complexes may ‘float as buoys’ at domain surfaces, whereas smaller TFs are less affected by the steric hindrance of dense chromatin (*27*).

### CDs provide an axis of control over DNA-DNA interactions

We show cohesin activity, while essential for TADs in ensemble Hi-C, to be dispensable for maintaining an interphase chain of consecutive CD. This is in line with a recent study by Bintu et al. demonstrating that local 3D domains can form *de novo* in the absence of cohesin, i.e. in a sequence-unspecific manner (*20*). We would conclude that CDs are not TADs, in that CDs represent the naive physical manifestation of chromatin derived primarily from its inherent polymer properties. As such, CD formation is independent of sequence specificity, and so cohesin-independent.

We postulate that the role of cohesin-CTCF is to add a layer of sequence-specificity on top of CDs exploiting the biophysically emergent structures to achieve physiologically relevant regulation between long-range *cis* elements (Fig. 6C, left). Without this cohesin-CTCF bridge between biology and physics two outcomes become apparent. CDs would be composed of non-physiologically relevant sequences, leading to an aberrant transcriptome (Fig. 6C, right). CD composition would also be highly variable from cell to cell leading to an erasure of TAD signal from ensemble population Hi-C. Both of these are known outcomes of cohesin ablation in interphase.

In contrast to cohesin, we have demonstrated that CDs and their nanoscale zonation can be locally and temporarily disrupted by active replication. The local relaxation of otherwise stable nucleosome aggregates may provide a window of opportunity for gene reactivation or for access by repair machineries to fix DNA damage from replication stress. In such scenarios the ATP-driven activity of cohesin as a loop extruder or as a sister-chromatid clamp may become critical, such as by stabilizing transient local higher-order chromatin structures that increase the fidelity of DNA double strand repair (*40*).

### Possible mechanisms of CD formation

The concept of an IC compartment permeating chromosome territories was introduced almost two decades ago (*4*). In such a model, the IC is a channel network that serves as transport highways for macromolecules to and from the nuclear pores. It was postulated that the PR together with the IC constitute the active nuclear compartment (*31*, *55*). Here by high-content analysis from 3D-SIM complemented by FIB-SEM, we show the IC to be a larger volume where a reticular network of CDs is found (Fig. S5B, C). Furthermore, transcriptional activity is enriched at a narrow CD chain-IC interface of < 60 nm in width (Fig. 3B). Most of the IC volume is therefore not involved in active transcription.

Co-staining for bulk RNA and DNA shows clear spatial exclusion of CDs from RNA-filled volumes. In fact, RNA transcripts, of which the vast majority are non-coding (*56*), are typically bound by intrinsically disordered proteins exerting multivalent interactions, known to have phase separating properties (*57*), and many transcription factors are known to have intrinsically disordered domains facilitating phase separation (*58*). It is thus very likely that phase separation contributes to the observed CD architecture by establishing a separation of RNA/protein micro-compartments, away from DNA-rich volumes.

Another driver of CD chain organization could be the default tendency of nucleosomes to self-interact into ever-larger collapsed polymer aggregates (‘melts’) (*59*). Chromatin domains with coherent dynamics have been reported in recent live-cell and correlated super-resolution imaging of replication domains (*60*). In support of these observations, our FIB-SEM shows CDs as clearly separated entities, many of which are tethered to each other via apparent single nucleosome wide linker filaments (Fig. 1B, arrows). These small linkers may offer mechanical coupling along the chain of CDs but remain flexible enough that each CD may move locally and independently from its neighbor. Being accessible, they also offer a site for the start of replication and transcriptional initiation.

These linkers also suggest that ‘stabilizing’ forces making CDs are counteracted by other forces. If this were not the case, both phase separation or polymer melts would eventually lead to a complete separation of chromatin into a single large unit. We have shown that chromatin destabilization can result from replication in S phase. The destabilization by DNA polymerase may provide a window for transcription factors to access previously inaccessible genes (Fig. 6B, right). We suggest that transcription and co-transcriptional processes supported by repulsive electrostatic forces (exerted by acetylated histone tails) are likewise involved in interphase (e.g.: at linker sites), tipping the local balance towards destabilization.

In conclusion, our novel nano-scale zonation model at the size of single physical CD entities allows for two independent axes of control. The first axis regulates DNA-RNA/protein interactions (Fig. 6B), the second regulates DNA-DNA *cis* contacts (Fig. 6C). In the first axis, the control of genome function can be achieved through changes in the CD chain surface area. The shape of CDs varies the quantity of chromatin available as substrate to epigenetic remodeler and transcription complexes at the CD-IC interphase. We show this to be true in euchromatic ‘A’ and heterochromatic ‘B’ compartments. In the second axis, CDs provide a structure that can be exploited by cohesin-based sequence-specific mechanisms for regulating interaction frequencies between long-range *cis* elements.

Based on these axes, our model predicts smaller domains with a larger surface-to-volume-ratio in constantly replicating pluripotent stem cells, and larger domains with a much-reduced surface-to-volume ratio in terminally differentiated or senescent cells that stop dividing. This would ensure genome plasticity, or structurally ‘lock-in’ and stabilize specific functional states, respectively. Specific DNA-protein interactions could be tested in defined subsets of CDs by immunofluorescence relative to single-TAD RASER-FISH. Regulation of DNA-DNA interactions dependent on CD activity states could be monitored by live-cell tracking of multiple *cis* loci within or between individual CDs.

## Acknowledgments

We thank Edith Heard for providing FISH probes against X-chromosomal TADs, Ron Schwessinger for help visualizing the C127 genome tracks, Afaf El-Sagheer and Arun Shivalingam for assistance in oligonucleotide synthesis, Anna Kreshuk and Jakop Pospisil for assistance with Ilastik and Imaris, as well as Phoebe Oldach and Eva Parisi for technical help. We are also indebted to Gleb Shtengel, Amalia Pasolli and Aubrey Weigel for help with FIB-SEM sample preparation and imaging, and David Brown and Rob Klose for providing H3K4me3 ChIP-Seq and ATAC-Seq data. We further wish to thank all colleagues who proofread and commented on the manuscript, in particular Fena Ochs, Justin Demmerle, Jiri Lukas, Claudia Lukas, Tim Nott and Rob Klose for valuable suggestions.

## Author contributions

E.M. and L.S. conceived the project and designed the experiments. C.S.X and H.F.H. conceived and conducted cryo-sample preparation and FIB-SEM imaging. E.M., R.O., H.A.S. and D.M.S.P. wrote the software and performed analyses. J.B. and V.B. devised the FISH protocol. E.M., R.O., J.B., A.R.C.F., S.d.O., C.I., J.R. and L.S. carried out experiments. E.M., R.O. and L.S. wrote the manuscript and generated the figures with contributions from all authors.

## Data and code availability

All data, parameters and scripts used are available for reproducing the results of this investigation. Raw and reconstructed 3D-SIM images are available at the Image Data Resource (IDR). The programs required to run all scripts for ChaiN are: ImageJ (FIJI distribution with SIMcheck), R and Octave. The live cell analysis via DFCC and Hi-D are run on Matlab with Image Processing, Signal Processing and Statistics and Machine Learning Toolboxes in a Windows environment. Scripts for ChaiN, DFCC and Hi-D analysis are available in the following GitHub repositories: https://github.com/ezemiron/Chain, https://github.com/romanbarth/DFCC, and https://github.com/romanbarth/Hi-D, respectively.

