## Supplementary Information for "Chromatin arranges in chains of mesoscale domains with nanoscale functional topography independent of cohesin"

**The PDF file includes:**
Materials and Methods
Figures S1-S9
Tables S1-S3
Captions for Movies S1-S6

**Other Supplementary Materials for this manuscript include the following:**
Movies S1 to S6

### Materials and Methods

#### *Plasmids*

pSpCas9(BB)–2A-Puro V2.0 targeting RAD21 (PX459 RAD21 (Hs) was described previously (*61*). A poly Glycine-Serine linker, Blasticidin resistance gene (BSD), GSG-P2A, mini auxin-inducible degron (mAID) and HaloTag were cloned between the KpnI and SalI site of a pUC19 vector by Gibson Assembly (NEB #cat E2611). The pUC19 CT-BSD_GSG_P2A-mAID-HaloTag was cloned between two 1 kb homology arms at the 3' of RAD21 to generate pUC19 RAD21 CT-BSD-GSG-P2A-mAID-HaloTag.

#### *BACs*

The BACs used for FISH labelling of a single TAD were a gift from E. Heard (*22*). They correspond to X chromosome "TAD H" characterized between genomic loci 102325818 and 102998976 (673158 bp). The four BACs required to cover this region are: 6-RP24-217I10 (102302874:102459532), 7-RP23-469A2 (102465652:102651023), 8-RP23-331L13 (102648562:102834423) and 9-RP24-396M14 (102804354:102976347) spanning a total length of 673473 bp. The 315bp difference between BACs coverage and TAD boundary calling is considered negligible over the 673kb total (<0.05% error). BAC probes were directly labelled by nick translation using CF-594 (Biotium).

#### *Cell culture*

Mouse mammary gland C127 and human cervical cancer HeLa H2B-GFP cells were grown in Dulbecco's Modified Eagle Medium (DMEM) supplemented with 10% FBS, 1% penicillin and streptomycin. Human colon carcinoma HCT116 Tet-OsTIR1 cells were cultivated in McCoy's 5A medium supplemented with 10% FBS, 2 mM L glutamine and 1% penicillin and streptomycin. Human lung fibroblast IMR-90 cells were grown in Dulbecco's Modified Eagle Medium (DMEM) supplemented with 10% FBS, 1% penicillin and streptomycin. Cells were incubated at 37°C, 5% $CO_2$ in a humidified incubator. At 90% confluence cells were trypsinized with 0.05-0.25% Trypsin in PBS or 1x trypsin replacement solution (TrypLE express, Gibco) and passaged to a new culture dish at appropriate dilutions (1:5-1:12 every 2-3 days).

#### *Generation of inducible RAD21 degradation system*

To generate a stable HCT116 Tet-OsTIR1 RAD21-mAID-Halo cell line, the pX459 RAD21 (Hs) plasmid and pUC19 RAD21 CT-BSD-GSG-P2A-mAID-HaloTag were transfected using FuGene HD (Promega #cat PRE2311). Two days post transfection, cells were grown in blasticidin-containing McCoy's 5A medium (5 $\mu$g/mL). After 10 days, single colonies were transferred to 96-well plates and homozygous clones were screened by PCR with primers listed in Table S2.

#### *EdU, EU and BrUTP labelling and perturbation experiments*

To identify cells in S-phase 10 $\mu$M 5-ethenyl-deoxyuridine (EdU) was added 15-30 min before fixation. G1 cells were identified by being negative for EdU pulse labelling and having a smaller nuclear size compared to G2 cells. For bulk RNA labelling 1 mM EU was added 60-90 min before fixation. To label nascent RNA, a 10 min BrUTP pulse labelling was performed by a scratch transcription labelling approach as previously described (*62*). For auxin-induced RAD21-degradation, HCT116 cells were cultured in the presence of 5 μg/mL doxycycline (Sigma, D9891) and 500 μM auxin (Sigma, cat# I5148) for 16 h and 6 h, respectively.

***RASER-FISH and probe labelling***

RASER-FISH maintains nuclear fine-scale structure by replacing heat denaturation with Exonuclease III digestion of one of the two DNA strands after UV-induced generation of nicks, and is suitable for high- and super-resolution imaging analysis (*39*). Briefly, cells were labelled overnight with a BrdU/BrdC mix (3:1) at a final concentration of 10 μM. Cells were fixed in 2% formaldehyde for 15 min and permeabilized in 0.2% Triton X-100/PBS (vol./vol.) for 10 min. Cells were incubated with 4',6-diamidino-2-phenylindole (DAPI; 0.5 μg/mL in PBS), exposed to 254 nm wavelength UV light for 15 min, then treated with Exonuclease III (NEB) at a final concentration of 5 U/μL at 37°C for 15 min. Labelled probes (100 ng each) were denatured in hybridization mix at 90°C for 5 min and pre-annealed at 37°C for 20 min. Coverslips were hybridized with prepared probes at 42°C.

***Immunofluorescence (IF) labelling***

IF labelling was performed as described in detail (*63*, *64*). The antibodies used in this investigation are listed in Table S3. One day before labelling, cells were grown on 22x22 mm #1.5H high precision 170 µm ± 5 µm coverslips (Marienfeld Superior). HCT116 Tet-OsTIR1 RAD21-mAID-Halo cells were fluorescently tagged by incubating with 50 nM HaloTag diAcFAM (Promega, cat# G8272) or 500 nM HaloTag JF488 for 30 min. HCT116 cells were incubated in HaloTag-free medium for 30 min to remove residual ligand. Prior to fixation, cells were washed twice with PBS. Cells were fixed in 2% formaldehyde/PBS (Sigma, #cat F8775) for 10 min, washed with 0.02% Tween-20/PBS (PBST) and permeabilized in 0.1-0.5% Triton-X-100/PBS. Coverslips were washed three times with PBST and incubated for 30 min in MaxBlock (Active Motif, cat# 15252). Cells were stained with primary antibodies against the protein of interest, washed three times with PBST and stained with fluorescently-labelled secondary antibodies. After washing, cells were post-fixed in 4% formaldehyde/PBS for 10 min and counterstained using 1-2 mg/μL DAPI or 1 μM SYTOX Green for 10 min. DAPI has a bias towards adenine and thymine over guanine and cytosine making AT-enriched satellite repeats near centromeric constitutive heterochromatin more intensively stained. This is particularly prominent in the chromocenters of mouse cells. This non-linear correlation between signal intensity and DNA concentration meant DAPI was replaced by SYTOX for all experiments possible. SYTOX has a weak affinity to bind RNA at high concentrations. Incubation with 1–10 U RNAse I/PBS prior to SYTOX counterstaining was performed as described (*62*). Coverslips were mounted in non-hardening Vectashield (Vector Laboratories, cat# H-1000) and stored at 4°C.

***Super-resolution image acquisition, reconstruction and quality control***

3D-SIM images were acquired with a DeltaVision OMX V3 Blaze system (GE Healthcare) equipped with a 60x/1.42 NA PlanApo oil immersion objective (Olympus), pco.edge 5.5 sCMOS cameras (PCO) and 405, 488, 593 and 640 nm lasers. For fixed cell imaging, 3D image stacks were acquired over the whole cell volume in z and with 15 raw images per plane (five phases, three angles). Spherical aberration was minimized using immersion oil with RI 1.514 for sample acquisition. The raw data was computationally reconstructed with

SoftWoRx 6.5.2 (GE Healthcare) using channel-specific OTFs recorded using immersion oil with RI 1.512, and Wiener filter settings between 0.002-0.006 to generate 3D stacks of 115 nm (488 nm) or 130 nm (592 nm) lateral and approximately 350 nm axial resolution. All SIM data was routinely and meticulously quality controlled for effective resolution and absence of artifacts using SIMcheck (*65*), an open-source Image J plugin to assess SIM image quality via modulation contrast-to-noise ratio, spherical aberration mismatch, reconstructed Fourier plot and reconstructed intensity histogram values. (for more details see Demmerle et al., 2017). Multi-channel acquisitions were aligned in 3D using Chromagnon software (*66*) based 3D-SIM acquisitions of multicolor EdU labelled C127 cells (*63*). 3D-SIM imaging of DAPI or SYTOX Green stained DNA typically provides a highly-contrasted spatial representation of chromatin distribution with a lateral resolution of wavelength-dependent down to ~100-115 nm in lateral and ~320 nm in axial direction (Fig. S1A).

For live cell 3D-SIM, HeLa H2B-GFP cells were seeded in a 35 mm μ-Dish, high Glass Bottom (Ibidi) using phenol red-free DMEM medium supplemented with 10 μM HEPES. Live cell imaging was performed at 37°C and 5% $CO_2$ supply by using an objective heater and a stage top incubator. We recorded for each time-point a central region over 7 z-positions with 125 nm distance between positions, covering a total range of 0.75 μm and totaling 105 raw images (5 phases x 3 angles x 7 z-positions) per timepoint. Using a 10 ms exposure time, frame rates of down to 0.5 Hz could be achieved, and typically 12 to 18 time points could be recorded until the photon budget was exhausted due to photobleaching. The reconstructed z-sections were processed in FIJI applying a maximum intensity z-projection followed by Bleach Correction (histogram matching) and Linear Stack Alignment with SWIFT (affine transformation) to compensate for photobleaching and nuclear motion/deformation.

***Quality validation***

3D-SIM imaging and subsequent quantitative analyses is susceptible to artefacts (*64*). For instance, bulk labelling of densely packed chromatin inside mammalian nuclei of several μm depth entails high levels of out-of-focus blur, which reduces the contrast of illumination stripe modulation and thus the ability to recover high frequency (i.e., super-resolution) structural information. We therefore carefully assessed and optimized system performance as well as raw and reconstructed data quality using the SIMcheck ImageJ plugin (*65*) (Fig. S1A). To exclude potential false positive calls, we used the modulation contrast-to-noise ratio (MCNR) map function of SIMcheck, which generates a metric of local stripe contrast in different regions of the raw data and directly correlates this with the level of high-frequency information content in the reconstructed data(*64*). Only immunofluorescent spot signals whose underlying MCNR values exceed a stringent quality threshold were considered while localizations with low underlying MCNR values were discarded to exclude any SIM signal which falls below reconstruction confidence. This criterion was applied to all datasets before feeding these to the analysis pipeline (Fig. S9). We noted that most nuclear spots (between a few hundred to several thousand per nucleus, depending on the target and the antibody) were in the size range of a 3D-SIM resolution-limited point signal, and that their intensities were similar to those few extranuclear spots originating from non-specifically bound primary-secondary antibody complexes containing fewer than 10 dye molecules. We therefore conclude that nuclear signals predominantly represent the binding of a primary-secondary antibody complex to a single epitope.

***Focused ion beam scanning electron microscopy (FIB-SEM)***
Cryo-fixed HeLa (CCL-2) cells were prepared for FIB-SEM using a previously described procedure (*36*). Briefly, cells grown on sapphire coverslips (3-mm diameter, 0.05-mm thickness; Nanjing Co-Energy Optical Crystal Co.) were subjected to high-pressure freezing using a Compact 01 high-pressure freezer (Wohlwend), followed by freeze-substitution (FS) under liquid N2 in FS medium (2% OsO4, 40 mM imidazole, 40 mM 1,2,4-triazole, 0.1% uranyl acetate, and 4% water in acetone) using an automated AFS2 machine (Leica Microsystems).

For resin embedding, samples were washed immediately after FS three times in anhydrous acetone for a total of 10 min, and embedded in Eponate 12 as previously described (*36*). FIB-SEM imaging was then performed using a customized Zeiss Gemini 500 crossbeam system previously described (*35*). The Zeiss Capella FIB column was repositioned at 90 degrees to the SEM column. The block face was imaged by a 240 pA electron beam with 1.0 keV landing energy at 200 kHz. The x-y pixel resolution was set at 4 nm. A focused Ga ion beam of 15 nA at 30 keV milled away 4 nm of material from the block face. The newly exposed surface was then imaged again. The milling / imaging cycle continued about once every 150 s for one month to capture the entire HeLa cell (50 x 8 x 68 $\mu m^3$). The raw image stack was aligned and registered using a Scale Invariant Feature Transform (SIFT) plug-in through Fiji. The 4 x 4 x 4 $nm^3$ (*67*) isotropic volume (16 x 3.2 x 24 $\mu m^3$) containing the nucleus was cropped from the whole cell stack for final analysis. Segmentation of FIB-SEM data was done employing machine learning using Ilastik 1.3.3 (*37*). 3D volume rendering after segmentation was performed with Imaris 9.2.

***ChaiN – a pipeline for high-content analysis of the 3D epigenome***
ChaiN (Chain high-throughput analysis of the *in situ* Nucleome) describes a pipeline of scripts for the automated high-throughput analyses in this investigation (Fig. S5). In brief, the counterstain/chromatin channel is used to generate (1) a nuclear mask and (2) segment chromatin topography into 7 intensity-based classes using an R script that is expanding on a Hidden Markov model (*68*). The modal intensity value (comprised of the large volume outside of the nucleus) is used to calibrate the relative intensity levels of each micrograph (due to unknown DNA dye incorporation and adjusted imaging conditions for optimized SIM imaging). Nucleoli were used as internal controls, as no signal should be detected in these volumes as expected. While this anchors the lower limit of the dynamic range, the upper limit is always relative to the maximum intensity value per micrograph, a limitation of the technique where cells of different conditions cannot be imaged together in the same micrograph for paired analysis. Nevertheless, based on the output of ChaiN, we can quantitatively assess multiple parameters, such as the overall volume of the PC+CD (combined classes 2-7, Fig. S5C) versus the IC (class 1), the PC-IC 3D surface, and the surface-to-volume-ratio. We note that the total number of discrete segmentation bins used in this analysis is arbitrary, with the sole criterion to group all non-detectable chromatin voxels as one class (class 1, IC).

The IF signals are thresholded by intensity using the Otsu algorithm, with all non-foci voxels replaced by zeros. The complement of this image is segmented by a 3D watershed algorithm and the 3D centroid coordinates are extracted from each segmented region's center of mass after weighting by local pixel intensities. To filter out potential false positive localizations, a local modulation-contrast-to-noise-ratio (MCNR) threshold is used to avoid potential artefacts skewing the statistical output (*64*, *65*) (Fig. S9). For each marker and condition at least 20 3D datasets from 2 biological replicates were acquired, each with many hundreds to several thousands of annotated IF foci.

The quality controlled and filtered centroid positions were then related to the chromatin-interchromatin volume in two ways: (1) The spot positions falling into each of the 7 segmented classes were counted, normalized to the respective class size, and their relative enrichment (or depletion) displayed in a heat map on a log 2-fold scale. Indexing IF coordinates on the segmented chromatin yields their enrichment or depletion profiles at different chromatin densities normalized to the volumes of each chromatin class.
(2) Segmentation of the 3D surface between class 1 and all other classes allows for a Euclidian distance calculation between the IF coordinates and the nearest CD chain surface voxel.

***Quantitative evaluation of chromatin dynamics***
Spatial correlation dynamics of chromatin was estimated by using Dense Flow reConstruction and Correlation (DFCC) (*32*). Briefly, DFCC was applied to estimate the 2D apparent motion of two consecutive images in an image sequence (N-1); the direction and magnitude of flow fields of each pixel (size = 41 nm) were estimated for H2B-GFP over a 20-image series and with a time interval of 2 s. Then, spatial auto-correlation of scalar fields representing motion direction was calculated by Fast Fourier Transforms given by

$$r(\Delta x, \Delta y) = \frac{\mathcal{F}^{-1}[\mathcal{F}(\gamma) \cdot \mathcal{F}^{*}(\gamma)]}{\langle\gamma\rangle\langle\gamma\rangle},$$

where $\mathcal{F}(\cdot)$, $\mathcal{F}^{-1}(\cdot)$, and $\mathcal{F}^{*}(\cdot)$ are the Fourier transformation, inverse Fourier transformation and the complex conjugate of the Fourier transformation, respectively. Correlation was calculated as a function of space lag and fitted using the Whittle-Màtern model correlation model to extract the correlation length and smoothness parameters. Correlation length ρc and the smoothness parameter v were derived from the regression and shown over the time lag. The parameters were averaged for each time interval over all accessible time points.

The Hi-D method was applied to estimate the biophysical properties of chromatin motion in living Hela cells at single-pixel resolution (41 nm) (*69*). Mean square displacement (MSD) curves were plotted over time lags representing the trajectories of chromatin in the entire nucleus. Then, a Bayesian inference approach was applied to test the following five principle models for each MSD curve in order to classify types of motion. Fitting of the MSD curve derived from trajectories can be expressed for anomalous diffusion (α), free diffusion constant (D), directed motion (V), or a combination thereof. The value of each biophysical parameter extracted by this fitting is then mapped in a 2D color map, and the five principal models are shown as a colored map directly on the nucleus (Fig. S3C). The five principle models for each MSD curve are

$$MSD_D(\tau) = 4D\tau + o$$
$$MSD_{DA}(\tau) = 4D\tau^{\alpha} + o$$
$$MSD_V(\tau) = v^2\tau^2 + o$$
$$MSD_{DV}(\tau) = 4D\tau + v^2\tau^2 + o$$

$$MSD_{DAV}(\tau) = 4D\tau^{\alpha} + v^2\tau^2 + o$$

where: D: Free diffusion; DA: Anomalous diffusion; V: Drift velocity of flow fields; DV: Free diffusion + drift; DAV Anomalous diffusion + drift; $\tau$: Time interval.

### Supplementary Figures

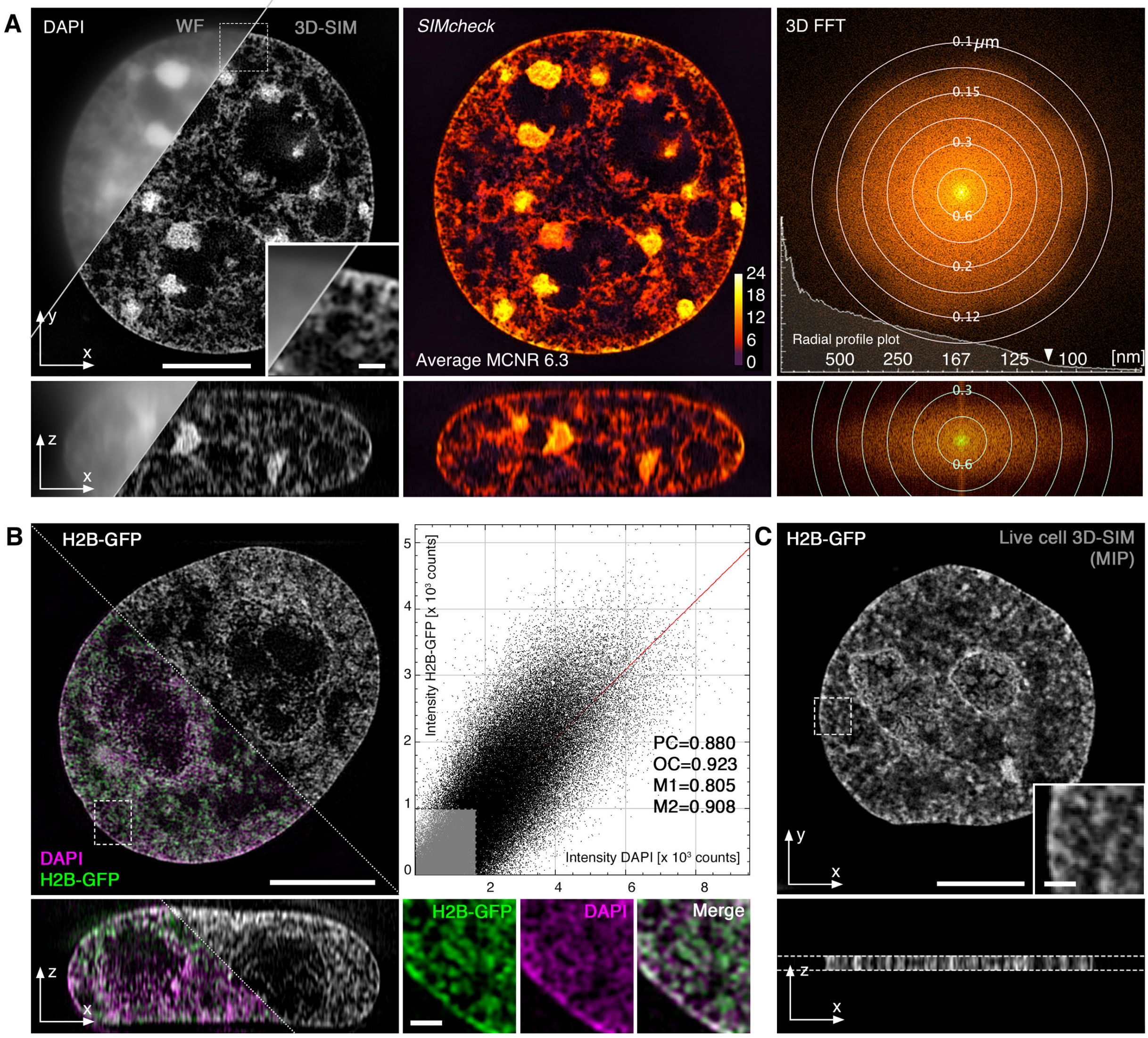


**Fig. S1. Super-resolution imaging of chromatin.**

(**A**) 3D-SIM data quality control. Left panel: Comparison of widefield (WF) and corresponding 3D-SIM image of a mouse C127 cell nucleus stained with DAPI (excited with 405 nm, detected in green emission of 500-550 nm); lateral (top) and orthogonal (bottom) cross sections are displayed. Middle and right panels: corresponding quality control analysis using SIMcheck (*65*), showing a mapping of local stripe modulation contrast (Modulation Contrast to Noise Ratio, MCNR) on the reconstructed cross sections (middle), and the corresponding axial and lateral Fourier plots with corresponding spatial resolution indicated by concentric rings (right). The superimposed radial profile plot indicates an effective lateral resolution of ~110 nm (arrow head). (**B**) 3D-SIM image of DAPI-stained (magenta) fixed H2B-GFP (green) cells and corresponding colocalization analysis showing strong correlation of both signals. PC: Pearson correlation coefficient; OC: Overlap coefficient; M1/M2: Manders' correlation coefficients DAPI vs. H2B-GFP and H2B-GFP vs. DAPI, respectively (using Otsu auto-threshold values indicated by grey box). (**C**) Live cell 3D-SIM image of histone H2B-

GFP in stably expressing HeLa cell (corresponding to Movie S2). To increase temporal resolution and reduce bleaching, only 7 z-planes covering ~1 $\mu$m depth are acquired and, after processing, displayed as maximum intensity projection (MIP), at the expense of compromising axial resolution (orthogonal view).

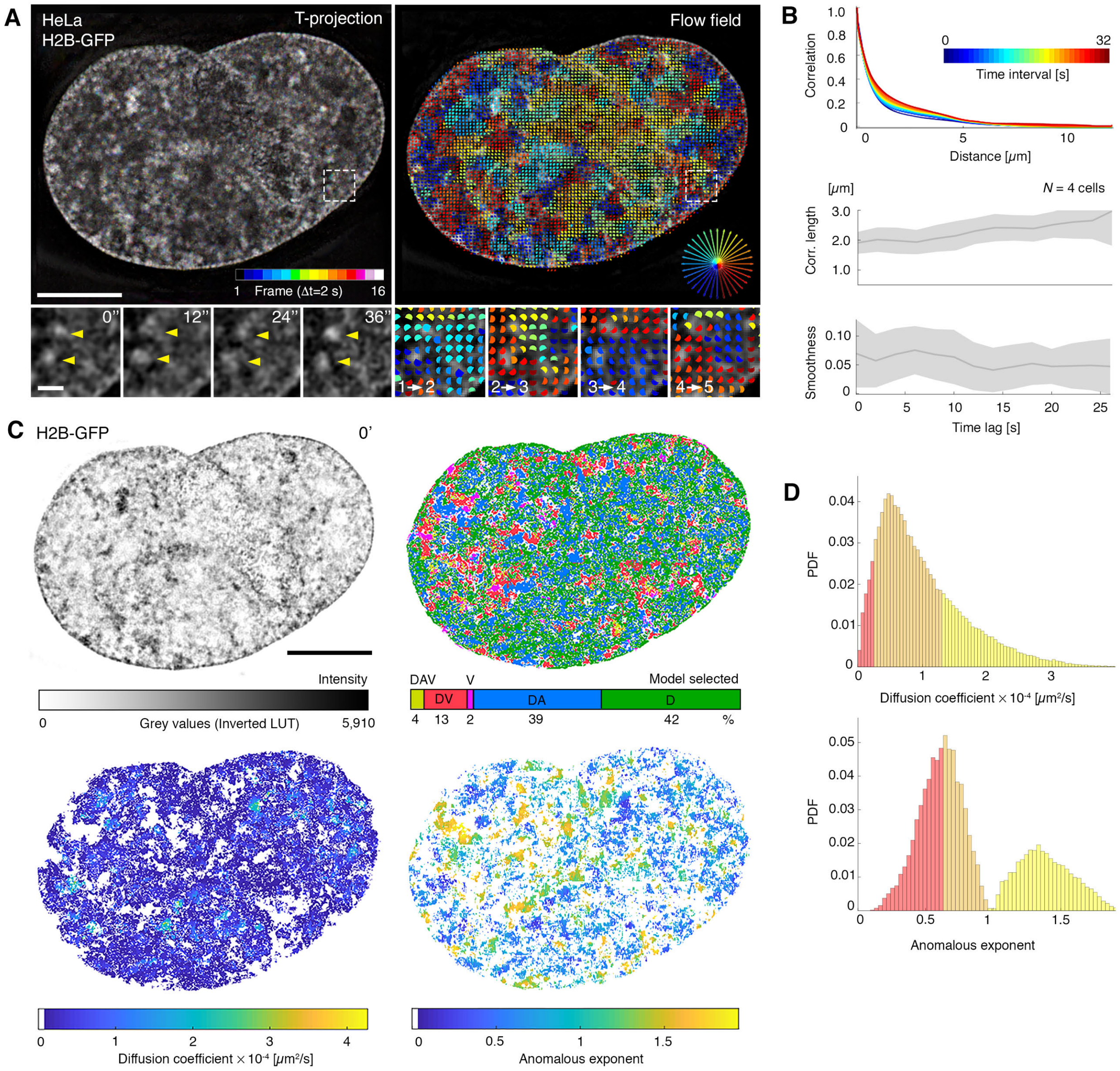


**Fig. S2. Quantitative analysis of chromatin dynamics demonstrates the coherent movement of nanodomains and heterogeneity of chromatin motion with sub-diffusive and super-diffusive dynamic regimes.**

(**A**) Live cell 3D-SIM of human HeLa cells stably expressing histone H2B-GFP recorded with intervals of 2 s (see Movie S1 for entire time series; representative time lapse recordings with 3 s and 30 s are shown in Movies S2 and Movie S3, respectively). Left panel: Color-coded projection of 16 consecutive time-points. Absence of color information indicates relative stability of higher order structures over time. Inset magnifications show selected time-points of the boxed region as a maximum projection of 7 z-sections covering ~1 $\mu$m depth. Inset magnifications of selected individual time-points showing coherently moving domains (arrow heads). Right panel: Overlay of flow field indicating motion direction for the first time interval. Fields are color-coded according to the direction of the displacement (Movie S4). Insets show analyses for the first 4 intervals with a time lag of 1 frame (Δ=2 s). Scale bars: 5 $\mu$m and 0.5 $\mu$m (insets). (**B**) Correlation function (top) calculated from dataset

shown in panel A as a function of distance at every accessible time interval (Δt is color coded from short to long time intervals, from blue to red, respectively) within the image series, each of them fitted to the Whittle–Matérn model. Correlation length (middle) and the smoothness parameter (bottom) are calculated over time for both direction and magnitude of flow fields. Results show a correlation length of ~ 2 to 3 $\mu$m for time intervals between 2 s and >20 s, indicating a change in correlation length of 0.4-0.5 $\mu$m between adjacent coherently moving domains (middle), for directional correlation of flow fields. This is accompanied by smoother transitions between these adjacent domains (bottom).
(**C**) Quantitative analysis of motion fields is applied using Bayesian model selection to define the types of diffusion processes acting on the chromatin fiber (top left) at the local and global scales. The spatial distribution of the selected models for each pixel is shown as a color map (top right), where D: Free diffusion; DA: Anomalous diffusion; V: Drift velocity of flow fields; DV: Free diffusion + drift; DAV Anomalous diffusion + drift. The mapping of the diffusion coefficient and the anomalous exponent in the lower panels highlight the heterogeneity of chromatin motion. (**D**) Histograms of diffusion coefficient and anomalous exponent values (panel C, lower right) plotted against the probability density function (PDF) after deconvolution using a general mixture model. For diffusion coefficient distributions, three mobility population groups, slow, intermediate and fast (shown as red, orange and yellow, respectively) are identified. The anomalous exponent distribution reveals two sub-diffusive populations (red and orange) and a third population with a super-diffusive regime (yellow).

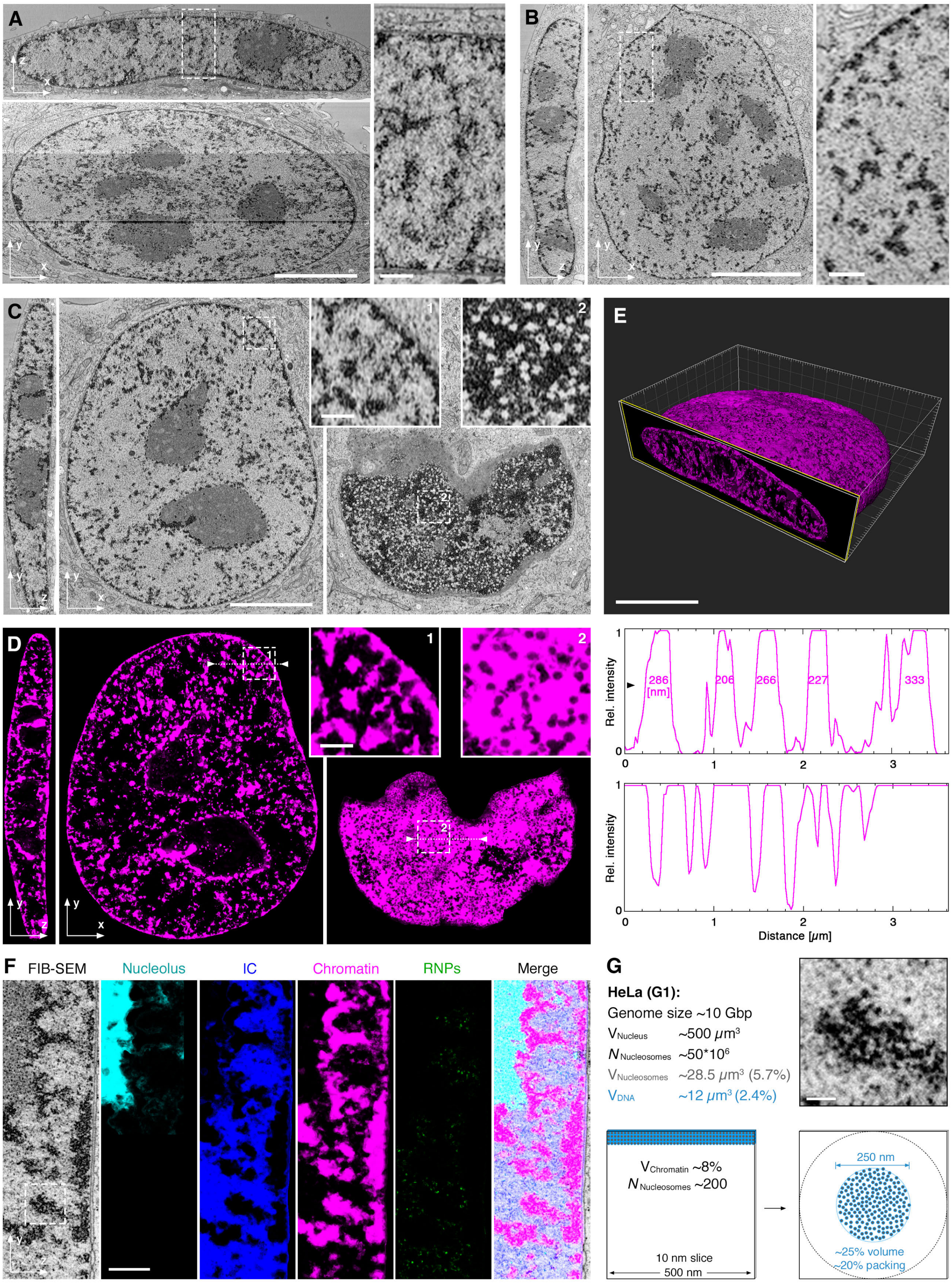
A
B
C
1
2
E
D
1
2
286 [nm]
206
266
227
333
Rel. intensity
Distance [μm]
F
FIB-SEM
Nucleolus
IC
Chromatin
RNPs
Merge
G
HeLa (G1):
Genome size ~10 Gbp
V Nucleus ~500 μm³
N Nucleosomes ~50*10⁶
V Nucleosomes ~28.5 μm³ (5.7%)
V DNA ~12 μm³ (2.4%)
V Chromatin ~8%
N Nucleosomes ~200
10 nm slice
500 nm
250 nm
~25% volume
~20% packing

**Fig. S3. Chains of CDs imaged with FIB-SEM.**

(**A**) Single FIB-SEM sections of a cryo-fixed HeLa cell, stained with osmium and uranyl acetate under cryo-conditions before resin embedding and imaging. A single 4 nm section was cut by a focused ion beam (FIB) in orthogonal direction to the growth surface and scanning electron microscopy (SEM) was recorded with 4 nm pixel size. Magnified regions display features in the axial dimension. (**B**) Single lateral and orthogonal FIB-SEM sections of a cryo-fixed U2-OS cell acquired with 8x8x8 nm voxel size. The magnified region displays features in the lateral dimension. Similar chromatin features can be observed for both HeLa and U2-OS nuclei. (**C**) Lateral sections of HeLa cell at the middle (left) and at the bottom (right) relative to growth surface. Data was downscaled to 20x20x20 nm voxel size for convenience. (**D**) Machine-learning-assisted segmentation of chromatin shown in magenta for the same data shown in panel C. The dimension of CDs is typically in the size range of 200-300 nm (line profile 1), with lamina associated CDs being no larger than CDs in the interior. Lamina associated CDs show no discernable linkers but instead form a continuous "melt" in an axis parallel to the nuclear lamina (right inset; line profile 2, dips in intensity correspond to nuclear pores). (**E**) 3D volume rendering of chromatin segmentation from the dataset in panel D. (**F**) Machine-learning-assisted segmentation of a high-resolution (4x4x4 nm voxel size) subvolume of the FIB-SEM dataset shown in Fig. 1E, F. Classification of 4 different states within the nuclear volume: the nucleolus (cyan), the interchromatin compartment (IC, blue), chromatin (magenta), and individual high contrast particles, likely RNPs (green). (**G**) Estimated average number, volume ratio and packing density of nucleosomes in a volume of 500x500x10 nm in a HeLa G1 cell given the genome size, the estimated number of nucleosomes, the volume of nucleosomes, and the volume of DNA. Distribution of the estimated number of 200 nucleosomes in an idealized CD of 250 nm diameter (bottom right) in comparison with a typical CD (boxed region in panel F) displayed at the same size scale (top right). Scale bars for all panels: 5 $\mu$m and 0.5 $\mu$m (for insets and magnified regions).

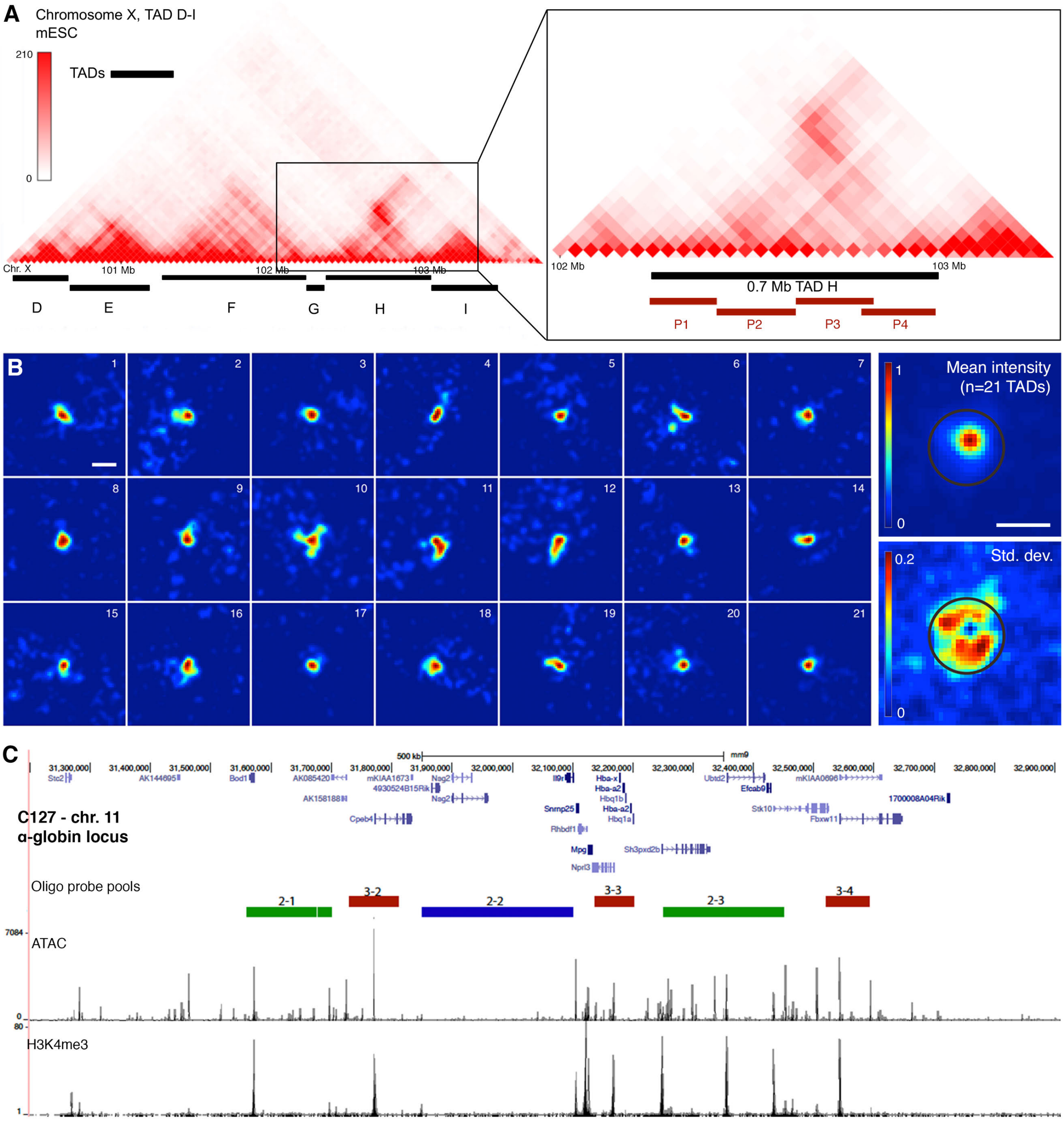


**Fig. S4. RASER-FISH for imaging chromatin conformation.**
(**A**) Hi-C heatmap from mouse ESCs (*70*) showing the interaction frequency around the 700kb TAD probe (probe H, red bars), (*22*) used in this study for RASER-FISH experiments. (**B**) Left: collection of TAD H signals on the X chromosome. Heatmaps of single z-planes are shown. The displayed z-planes contain the brightest pixel of the TAD z-stack. Signal intensity was normalized to the brightest pixel. Top right panel: quantification of the mean intensity of 21 TADs. For each TAD, the z-plane containing the brightest pixel was selected, centered with respect to this pixel and normalized to its signal intensity (top). Bottom right: the standard deviation (SD, bottom) describes the degree of variation in maximum elongation of TADs, indicated with a diameter of ~500 nm. Scale bar: 0.5 $\mu$m. (**C**) Chromatin features in C127 cells showing sequential domains (see Fig. 2C) characterized by the presence or absence of H3K4me3 ChIP-Seq and ATAC-Seq peaks.

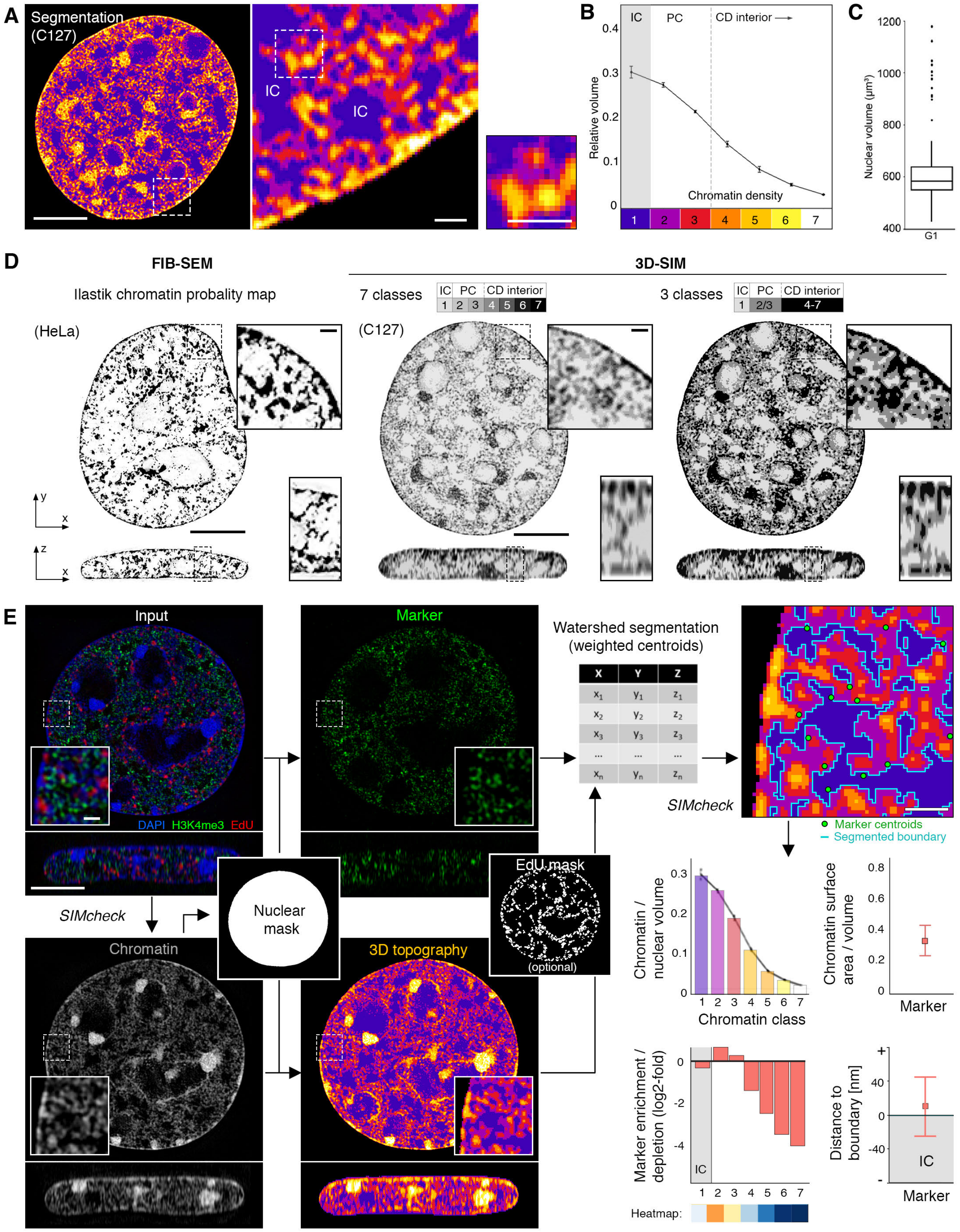


**Fig. S5. Systematic analysis of the 3D epigenome.**
(**A**) Segmentation of chromatin staining into 7 intensity classes (Methods) at three different serial magnifications. (**B**) Quantification of class sizes from interchromatin (IC, class 1), perichromatin (PC, class 2-3) and core regions of CD chains (CD interior, classes 4-7) shows a decrease of nuclear volume fractions for higher classes. The IC and PC together account for ~75% of nuclear volume from SIM data. Error bars indicate 95% CI. (**C**) Box plot

of nuclear volumes in C127 G1 cells from all segmented nuclear voxels. $N = 433$ cells (panels B and C). (**D**) Comparison of segmentation methods for FIB-SEM (left) and 3D-SIM (center, right) data. Lateral (top) and orthogonal cross section (bottom) are shown. Grouping the 7-class segmentation (center) into class 1, classes 2-3 and classes 4-7 (right) most closely resembles the FIB-SEM segmentation (left), validating the clustering of classes into 3 main states: IC, PC and CD interior (as in panel B).) Note that the anisotropic resolution of 3D-SIM along the z-direction, highlighted in the orthogonal sections (bottom), skews the comparison with isotropic FIB-SEM data. (**E**) Schematic representation of the ChaiN analysis workflow. Inputs for the workflow are multi-channel 3D-SIM datasets that have been pre-processed, quality-controlled, thresholded and aligned. First, a nucleus mask is generated based on the chromatin channel. Secondly, within this mask, marker spots are segmented by intensity and their weighted centroid positions are determined. Simultaneously, the chromatin channel is segmented into 7 intensity classes as detailed above to describe the 3D topography of chromatin. Finally, metrics from both marker and chromatin channels are used to describe the 3D nucleome, such as the nuclear proportion of each chromatin class, the chromatin surface-to-volume ratio, the marker enrichment or depletion in each chromatin class, or the mean distance (± 95% CI) of markers to the segmented chromatin-interchromatin edge. Analysis is performed by linking scripts written in R and Octave. Scale bars for all panels: 5 $\mu$m and 0.5 $\mu$m (insets).

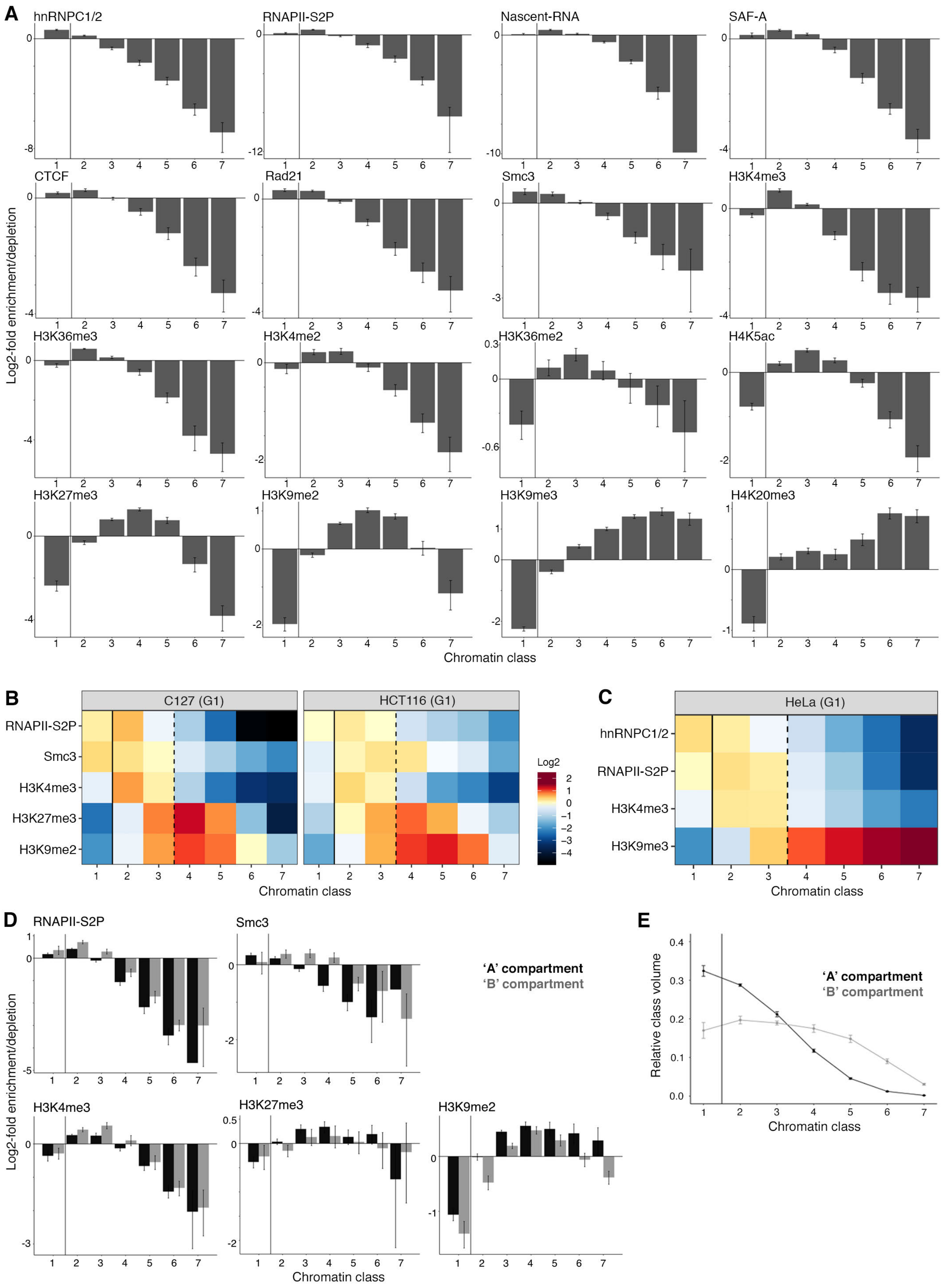
A
hnRNPC1/2
RNAPII-S2P
Nascent-RNA
SAF-A
CTCF
Rad21
Smc3
H3K4me3
H3K36me3
H3K4me2
H3K36me2
H4K5ac
H3K27me3
H3K9me2
H3K9me3
H4K20me3
Log2-fold enrichment/depletion
Chromatin class
B
C127 (G1)
HCT116 (G1)
RNAPII-S2P
Smc3
H3K4me3
H3K27me3
H3K9me2
Log2
Chromatin class
C
HeLa (G1)
hnRNPC1/2
RNAPII-S2P
H3K4me3
H3K9me3
Chromatin class
D
RNAPII-S2P
Smc3
H3K4me3
H3K27me3
H3K9me2
'A' compartment
'B' compartment
Log2-fold enrichment/depletion
Chromatin class
E
Relative class volume
'A' compartment
'B' compartment
Chromatin class

**Fig. S6. Functional marker distributions in whole mouse and human cell lines and macro-compartments.**

(**A**) Bar plots of the log-2 fold enrichment or depletion of each analyzed marker in each of the segmented chromatin classes, in mouse C127 G1 cells, relative to chromatin class volume. Numbers of cells for each marker is detailed in Table S1. (**B**-**C**) Representative selections of markers in mouse C127 cells show the same zonation patterns with the same antibodies in human HCT116 colon carcinoma cells (panel B) and HeLa cells (panel C), suggesting chromatin accessibility of PC (classes 2-3) vs CD interior (classes 4-7) as a universal regulator of epigenetic zonation. Number of cells: C127 = 170, HCT116 = 70, HeLa = 41. (**D**) Bar plots of the over/under representation of representative markers in each of the segmented G1 macro-scale A/B chromatin compartments, showing a relatively conserved zonation pattern between active and repressive markers. Plots show average log-2 value, error bars indicate 95% confidence interval. Number of cells = 170. (**E**) Composition of chromatin classes in either A or B regions. Plot shows relative class volume, error bars indicate 95% confidence interval. Number of cells = 170.

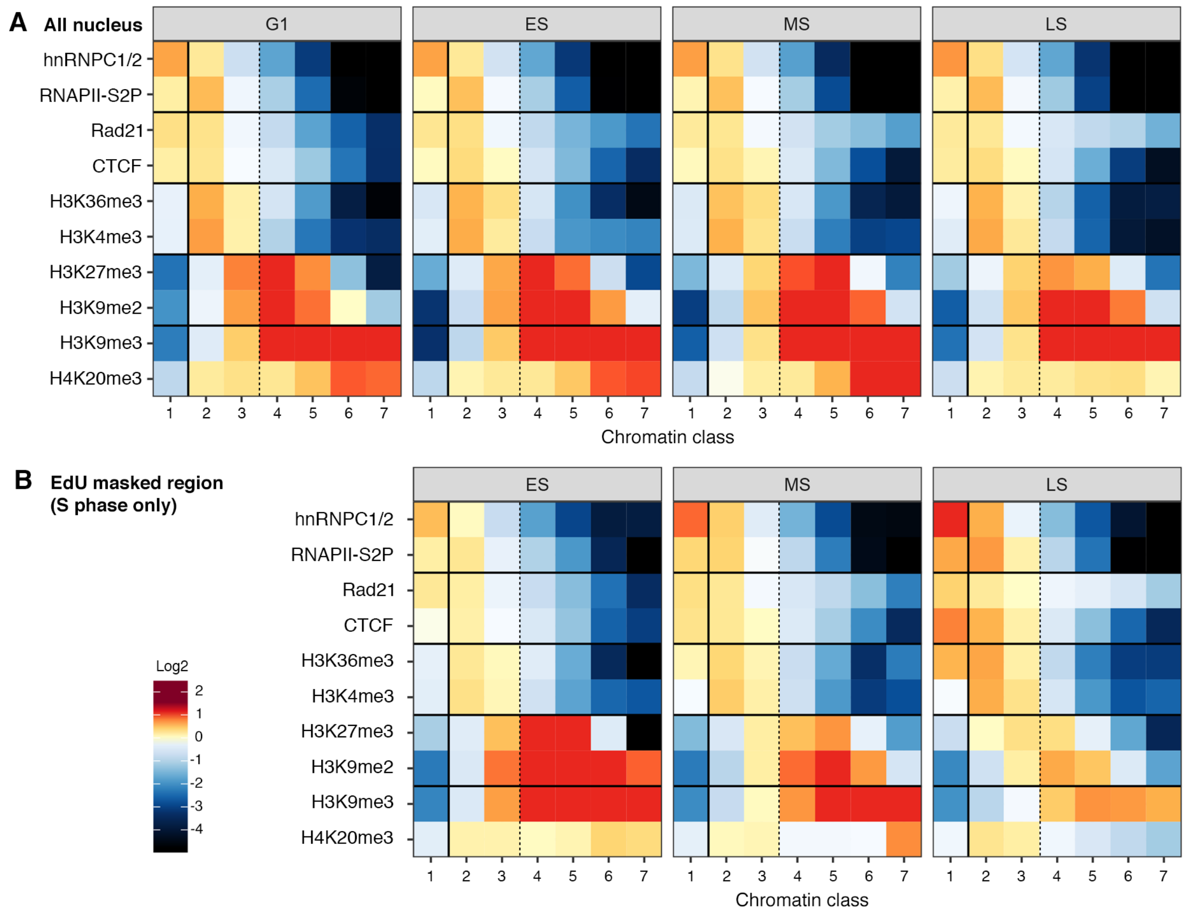


**Fig. S7. Functional marker distribution in different S-phase stages of mouse C127 mammary epithelial cells.**

(**A**-**B**) Heatmaps of enrichment or depletion of IF signals relative to a random distribution (plotted in log2-fold change), per chromatin class, in the whole nuclear volume (A) or for a local EdU-submask (B). One heatmap per cell cycle stage. Number of cells: G1 =320, ES = 225, MS = 227, LS = 230.

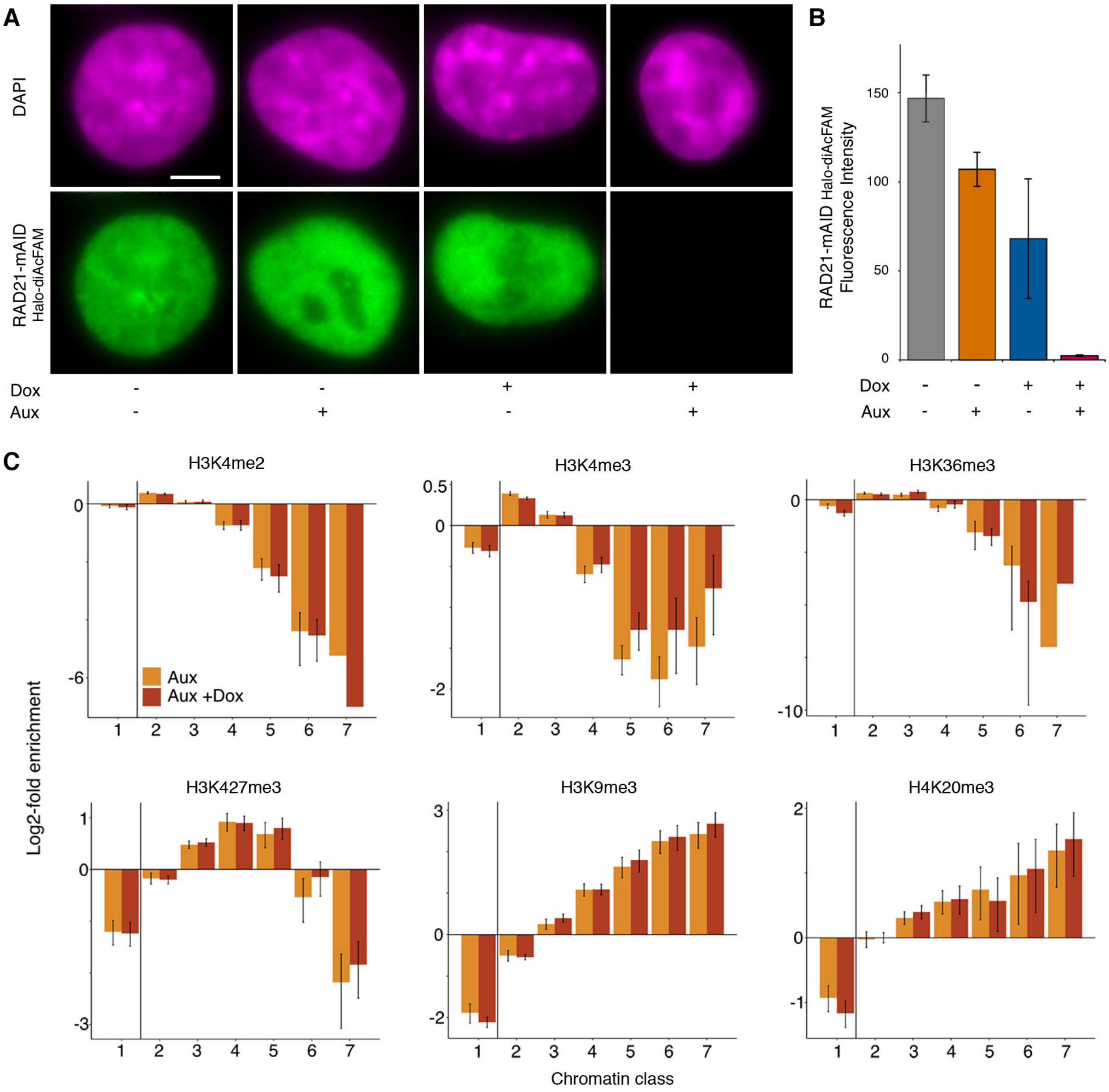


**Fig. S8. Characterization of HCT116 Tet-OsTIR1 RAD21-mAID-Halo cell line and functional marker distribution for RAD21 ablation.**

(**A**) Representative widefield fluorescent DAPI and RAD21-mAID signals after mock induction, 2 h auxin (aux), 16 h doxycycline (dox), or both 2 h aux and 16 h dox (left to right respectively). Scale bar: 5 $\mu$m. (**B**) Quantification of the fluorescence intensity in the different conditions of (A). N = 10 cells per condition (error bars = SD). (**C**) Bar plots of the over/under representation of each histone PTM analyzed in each of the segmented chromatin classes, with or without RAD21 ablation (6h aux +/- 16 h dox), in HCT116 cells. Plots show average log-2 value, error bars indicate 95% confidence interval. Numbers of cells for each marker is detailed in Table S1.

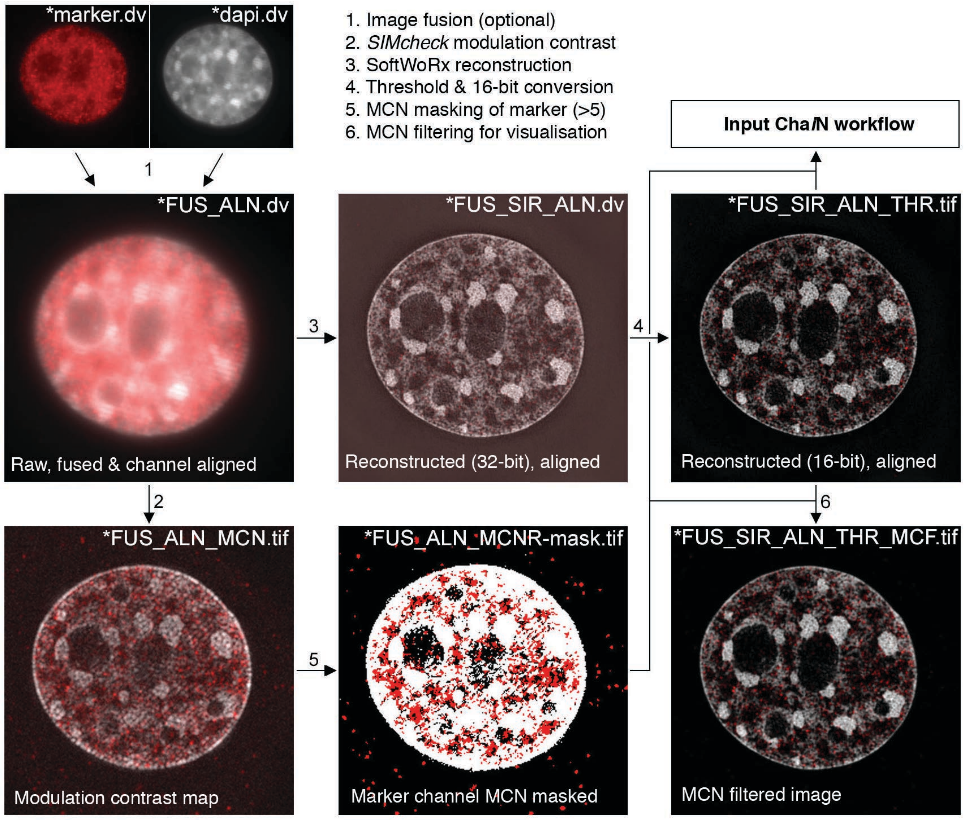


**Fig. S9. Schematic representation of image pre-processing prior to automated ChaiN pipeline.**
Both marker and chromatin signals were recorded using the same fluorescence channel, as capturing the bleed-through of the chromatin signal improved alignment quality checks using Chromagnon (*66*). Reconstructed images were thresholded and the modulation-contrast-to-noise (MCN) was calculated using SIMcheck (*65*). A MCNR-mask was generated (using quality cut-off at: MCN=5). Thresholded images together with the MCNR-mask were used as input for ChaiN to discard sub-quality voxels in the marker signal foci, reducing the effect of 3D-SIM artefacts on subsequent image analysis.

**Table S1. Average number of foci analyzed per marker.**

| | **C127** | | | |
|---|---|---|---|---|
| **Marker** | **G1** (*N*=433) | **ES** (*N*=225) | **MS** (*N*=227) | **LS** (*N*=230) |
| CTCF | 7130 ± 2025 (40) | 4784 ± 1699 (30) | 4052 ± 2732 (30) | 3878 ± 3360 (30) |
| H3K27me3 | 2590 ± 1500 (40) | 1906 ± 1727 (20) | 1940 ± 1734 (18) | 1949 ± 1576 (20) |
| H3K36me2 | 3260 ± 1993 (20) | - | - | - |
| H3K36me3 | 1323 ± 515 (40) | 2822 ± 1396 (30) | 2541 ± 1153 (30) | 392 ± 356 (30) |
| H3K4me2 | 2708 ± 1089 (20) | - | - | - |
| H3K4me3 | 5556 ± 1207 (40) | 3692 ± 1777 (30) | 2781 ± 2416 (29) | 3104 ± 2778 (30) |
| H3K9me2 | 4787 ± 988 (30) | 1169 ± 535 (15) | 2753 ± 1405 (20) | 2548 ± 1867 (20) |
| H3K9me3 | 1908 ± 1603 (30) | 688 ± 230 (20) | 809 ± 412 (20) | 664 ± 229 (20) |
| H4K20me3 | 4523 ± 356 (20) | 2598 ± 583 (20) | 1959 ± 1196 (20) | 2344 ± 392 (20) |
| H4K5ac | 5177 ± 1527 (20) | - | - | - |
| hnRNPC1/2 | 4625 ± 781 (20) | 1224 ± 423 (20) | 865 ± 174 (20) | 614 ± 220 (20) |
| Nascent RNA | 2797 ± 1124 (13) | - | - | - |
| RAD21 | 1272 ± 391 (20) | 981 ± 193 (10) | 991 ± 102 (10) | 1146 ± 244 (10) |
| RNAPII-S2P | 5027 ± 1319 (40) | 2830 ± 1860 (30) | 2698 ± 2530 (30) | 3019 ± 2360 (30) |
| SAF-A | 4655 ± 1821 (20) | - | - | - |
| Smc3 | 615 ± 263 (20) | - | - | - |

| | **HCT116 (RAD21-mAID)** | | **HeLa** |
|---|---|---|---|
| | **+ RAD21** (*N*=80) (6 h Auxin) | **- RAD21** (*N*=104) (16 h Dox + 6 h Auxin) | **G1** (*N*=41) |
| H3K27me3 | 1077 ± 516 (20) | 1379 ± 407 (30) | - |
| H3K36me3 | 551 ± 138 (10) | 790 ± 263 (12) | - |
| H3K4me2 | 2620 ± 413 (10) | 3063 ± 412 (12) | - |
| H3K4me3 | 7409 ± 1252 (20) | 8155 ± 1420 (30) | 7341 ± 1785 (10) |
| H3K9me3 | 1182 ± 204 (10) | 1771 ± 174 (10) | 5837 ± 525 (11) |
| H4K20me3 | 2316 ± 387 (10) | 2246 ± 272 (10) | - |
| hnRNPC1/2 | - | - | 7531 ± 1463 (10) |
| RNAPII-S2P | - | - | 3719 ± 819 (10) |

Average number of IF spots ± SD of each marker per cell for (*N*) number of cells in each condition, after high stringency filtering to avoid false positives. Cell cycle stages are abbreviated to ES, MS and LS for early, mid and late S phase. To maintain consistency, the same mammalian antibodies were used throughout this study, however the affinity of some antibodies is greater or decreased in either mouse (C127) or human (HCT116, HeLa) cells.

**Table S2. PCR primer sequences.**

| Name | Orientation | Sequence |
|---|---|---|
| B2A-BSD | Fw | GCTCCTATTAGCCCTCCCACACATAACC |
| | Rev | GGATGTATAAAGGTTCCGGAGCCACCAAC |
| RAD21 (frag. 3) | Fw | AGGGCTAATAAGGAGCTAGAAGCATTATAGC |
| | Rev | TTGCATGCCTGCAGGGTTCCAAACCAGGAGTGTG |
| RAD21 (frag. 1) | Fw | AATTCGAGCTCGGTACTGAATGTCTGCAAAATGC CAAGC |
| | Rev | CTTCTCCTTGCCGGAAATCTCGAGCGTC |
| mAID | Fw | TTTCCGGCAAGGAGAAGAGTGCTTGTC |
| | Rev | CCGGAACCTTTATACATCCTCAATCGATTTTC |
| Genomic primer | Fw | AGCATGAGATTGGAGGGGA |
| | Rev | GCCACTCAACATTATTACATGCA |

**Table S3. Primary and secondary antibodies used in this study.**

| Antibodies | Source | Identifier |
|---|---|---|
| *Primary antibodies:* | | |
| Rabbit mAb anti-hnRNP C1 + C2 | Abcam | ab133607 |
| Mouse mAb anti-Rad21 | Merck Millipore | 05-908 |
| Rabbit pAb anti-SMC3 | Bethyl Laboratories Inc. | A300-060A |
| Rabbit mAb anti-CTCF | Cell Signalling Tech. | D31H2 |
| Rabbit pAb anti-SAF-A | Abcam | ab20666 |
| Rabbit pAb anti-RNAPII ser2P | Abcam | ab5095 |
| Rabbit pAb anti-H3K36me2 | Active Motif | 39255 |
| Rabbit pAb anti-H3K36me3 | Active Motif | 61102 |
| Mouse mAb anti-H3K4me2 | Active Motif | 39679 |
| Rabbit pAb anti-H3K4me3 | Active Motif | 39159 |
| Mouse mAb anti-H3K27me3 | Abcam | ab6002 |
| Rabbit pAb anti-H4K20me3 | Abcam | ab9053 |
| Mouse mAb anti-H3K9me2 | Abcam | ab1220 |
| Mouse mAb anti-H3K9me3 | Active Motif | 61013 |
| Rabbit mAb anti-H4K5ac | Merck Millipore | 04118 |
| *Primary antibodies (directly conjugated):* | | |
| Alexa Fluor 488 Rat mAb anti-BrdU | Abcam | ab220074 |
| *Secondary antibodies:* | | |
| Alexa Fluor 488 Goat pAb anti-Mouse-IgG | Thermo Fisher Scientific | A11029 |
| Alexa Fluor 488 Goat pAb anti Rabbit-IgG | Thermo Fisher Scientific | A11034 |
| Alexa Fluor 594 Donkey pAb anti-Mouse-IgG | Thermo Fisher Scientific | A21203 |
| Alexa Fluor 594 Donkey pAb anti-Rabbit-IgG | Thermo Fisher Scientific | A21207 |

**Movie S1. Super-resolution imaging of chromatin dynamics.**
Live cell 3D-SIM of a human HeLa cell stably expressing histone H2B-GFP. Dataset corresponds to Figure 1D, recorded with time intervals of 2 s. For each frame, 7 z positions with a distance of 125 nm were acquired, totaling 105 raw images (5 phases, 3 angles, 7 z positions), covering a depth of 0.75 $\mu$m. The reconstructed z-sections were maximum intensity projected, corrected for bleaching, and registered to compensate for cell motion and nuclear deformation. For the movie an ‘Orange Hot’ LUT was chosen.

**Movie S2. Super-resolution imaging of chromatin dynamics.**
Live cell 3D-SIM of a human HeLa cell stably expressing histone H2B-GFP. Dataset corresponding to Figure S3A, recorded with time intervals of 3 s. Processing was performed as described for Movie S1.

**Movie S3. Super-resolution imaging of chromatin dynamics.**
Live cell 3D-SIM of a human HeLa cell stably expressing histone H2B-GFP. Dataset corresponding to Figure S3B, recorded with time intervals of 30 s. Processing was performed as described for Movie S1.

**Movie S4. Quantitative analysis of chromatin dynamics.**
Flow field analysis of the dataset shown in Movie S1, corresponding to Figure S3A, highlighting changes in motion direction between frames, characteristic for an elastic and coherent motion of chromatin features. Note that only changes for time lags of 1 frame are displayed, while the for subsequent correlation analyses (Figure S3B) all time lags are included.

**Movie S5. FIB-SEM sectioning through a HeLa cell nucleus.**
Serial sections were taken perpendicular to the growth surface with a 4 nm isotropic voxel size. Movie shows a representative subvolume of the dataset presented in Figures 1E, F and Figure S3C-E. Data was downscaled to 8 nm voxel size and jpeg compressed to reduce file size.

**Movie S6. 3D volume rendering of FIB-SEM imaged HeLa cell nucleus.**
Movie shows subvolume of the dataset presented in Figures 1E, F and S3C-E at 4 nm isotropic voxel resolution. Chromatin (magenta) and RNPs (green spots) were segmented using Ilastik software and rendered with Imaris.
